# Dispensing with unnecessary assumptions in population genetics analysis

**DOI:** 10.1101/2022.09.12.507656

**Authors:** Olivier Labayle Pabet, Kelsey Tetley-Campbell, Mark J. van der Laan, Chris P. Ponting, Sjoerd Viktor Beentjes, Ava Khamseh

**Author notes:** For correspondence (AK); (SVB). These authors contributed equally to this work (Sjoerd Viktor Beentjes, Ava Khamseh).

## Abstract

Parametric assumptions in population genetics analysis – including linearity, sources of population stratification and additivity of variance as part of a Gaussian noise – are often made, yet their (approximate) validity depends on variant and traits of interest, as well as genetic ancestry and population dependence structure of the sample cohort. We present a unified statistical workflow, called TarGene, for targeted estimation of effect sizes, as well as two-point and higher-order epistatic interactions of genomic variants on polygenic traits, which dispenses with these unnecessary assumptions. Our approach is founded on Targeted Learning, a framework for estimation that integrates mathematical statistics, machine learning and causal inference. TarGene maximises power whilst simultaneously maximising control over false discoveries by: (i) guaranteeing optimal bias-variance trade-off, (ii) taking into account potential covariate non-linearities, sources of population stratification and dependence structure, and (iii) detecting genetic non-linearities. The necessity of this model-independent approach is demonstrated via extensive simulations. We validate the effectiveness of our method by reproducing previously verified effect sizes on UK Biobank data, whilst simultaneously discovering non-linear effect sizes of additional allelic copies on trait or disease, in a PheWAS study involving 781 traits. Specifically, we demonstrate genetic non-linearity at the FTO locus is significant for 54 traits in this study. We further find three pairs of epistatic loci associated with skin color that have been previously reported to be associated with hair color. Finally, we illustrate how TarGene can be used to investigate higher-order interactions using three variants linked to the vitamin D receptor complex. TarGene provides a platform for comparative analyses across biobanks, or integration of multiple biobanks and heterogeneous populations to simultaneously increase power and control for type I errors, whilst taking into account population stratification and complex dependence structures.

## Introduction

Genome-wide association studies (GWAS) aim to quantify associations between DNA variants and complex traits or disease, given a sample population of interest which may contain individuals from various genetic ancestries or dependence. Over the past 15 years, many risk loci have been identified to be associated with disease or traits (***Visscher et al., 2017***), through a variety of statistical models such as logistic, linear and linear mixed models (LMMs) (***Uffelmann et al., 2021***) most of which rely on strong parametric assumptions. These assumptions include linearity of the DNA variant-trait relationship, linearity of the functional dependence of trait on covariates and confounders such as sources of population stratification and other fixed effects including batch, age and sex, as well as additivity of variances as part of a Gaussian noise encoded as random effects, in the commonly employed linear mixed models (LMMs) (***Uffelmann et al., 2021***; ***Svishcheva et al., 2012***; ***Lippert et al., 2011***). The validity of these assumptions depends on the variants and traits of interest, as well as the specific population structure of the sample cohort, such as genetic ancestry and kinship dependence amongst the individuals. If the assumptions lead to biased estimates of the quantities of interest, this bias is exacerbated by the sample size, i.e., the larger the sample size, the further away the estimates will get from the ground truth (Fig. 1). This Curse of Big Data is especially likely because modern biobank-scale data sets, with their ever-growing size of 10^5^ − 10^6^ individuals, result in very small variance, which exposes bias (Fig. 1A), and leads to overly optimistic p-values. There is therefore a need to develop robust statistical and machine learning methodologies that are mathematically guaranteed to maximise power and control over false discoveries simultaneously.

**Figure 1.**
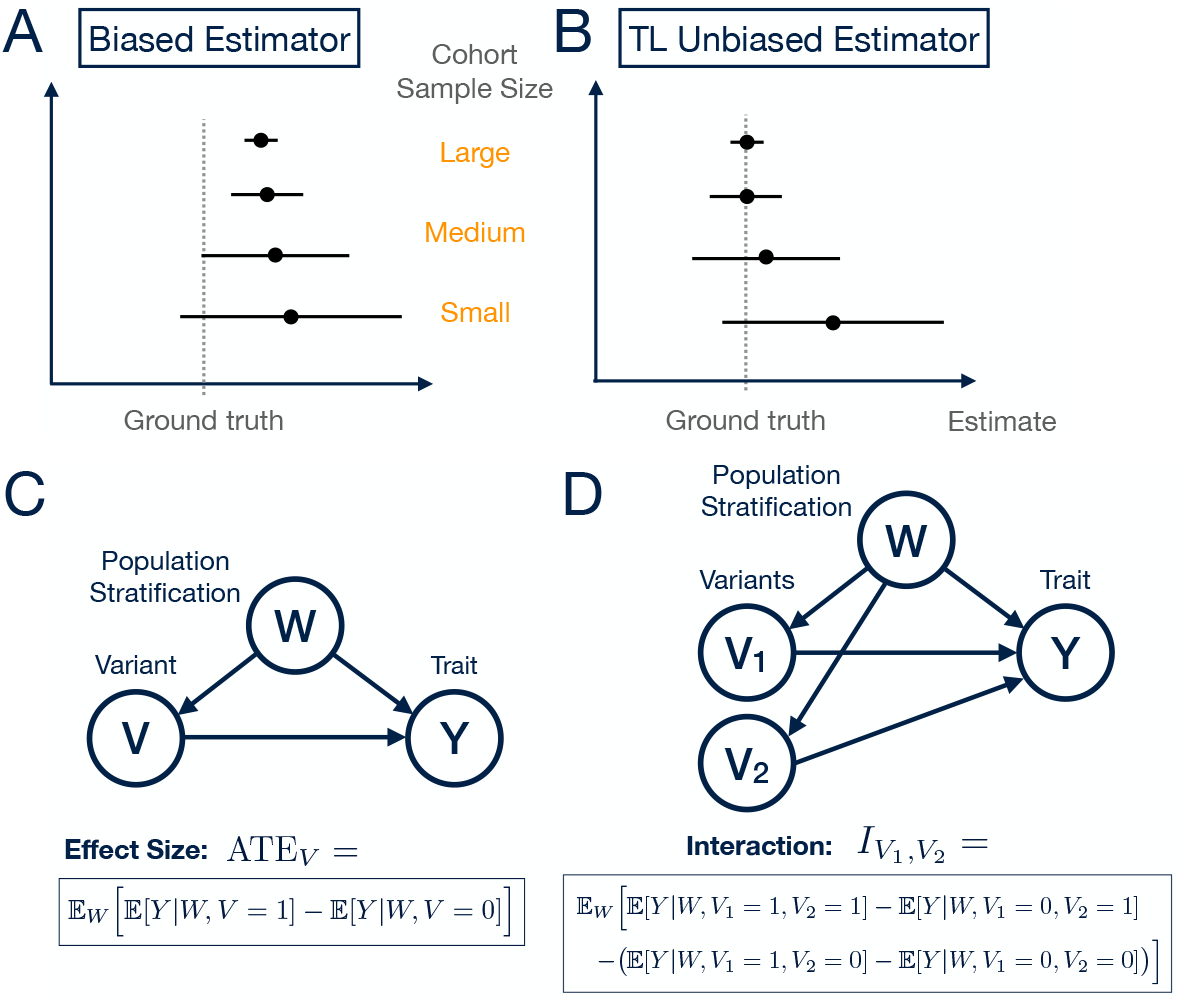
**A** and **B**: The Curse of Big Data. **A**: As sample size increases the bias of an estimator may not shrink sufficiently fast relative to the reduction in variance, leading to incorrect predictions for large sample sizes. Extensive simulations demonstrating this phenomenon are presented in Fig. 4 using different ground truth models fitted with a misspecified model, under various variant-covariate dependence structures (see Methods). **B**: Targeted Learning estimator provides correct statistical inference once it has been optimised for bias-variance trade-off in the TMLE step. **C**: Model-independent definition of effect size (Average Treatment Effect, ATE) of variant *V* on trait *Y*. We condition on sources of population stratification and other confounders, *W*, when estimating the ATE. We do this to get closer to a causal estimate of the variant on trait, up to linkage disequilibrium (LD). With additional molecular information to reduce LD and/or fine-mapping these estimates can approach being causal over associative. For a binary variable *V* the interpretation is: “Having correctly adjusted for confounders (sources of population stratification), what is the difference in the expected value of trait when a particular variant *V* is present (*V* = 1) as compared to when it is not (*V* = 0)?” Generalisation to a categorical variable *V* = 0, 1, 2 is trivial, as all combinations can be written similarly. **D**: Model-independent definition of 2-point interaction, an extension of ATE with more than one variant, has been further generalised to higher-order interactions among *n* variants (***Beentjes and Khamseh, 2020***). Interpretation: “Having adjusted for confounders, is the effect of variant *V*_1_ on trait *Y* modulated by the status of variant *V*_2_ and, if so, by how much and with which sign?” This defines an epistatic interaction between variants *V*_1_ and *V*_2_ with respect to a trait.

Rather than addressing the root cause of each bias, current methods often seek to ameliorate their effects post hoc (***Fusi et al., 2014***). An exception to this generalisation is KnockoffGWAS which makes no parametric assumptions regarding the distribution of the phenotype conditional on the genotypes (***Sesia et al., 2021***). This method also controls the false discovery rate (FDR) whilst accounting for population structure. Nevertheless, KnockoffGWAS does not compute effect sizes or epistatic interactions, and only reports statistical significance, and thus does not directly (i) detect genetic non-linearity, (ii) infer the strength and sign of variant effect sizes and interactions, or (iii) determine whether or not a variant of interest is stratified across the population. Estimation of these quantities is essential for explaining how variants, via biological mechanisms and regulatory functions, modify a trait or disease risk.

New Machine Learning (ML) approaches have begun to be applied to many aspects of genetics and genomics but not, until recently, to population genetics analysis. A recent ML method (***McCaw et al., 2022***) partially addresses the issue of non-linear and interacting covariates in the phenotype-genotype relations by modeling these using deep neural networks. However, this approach relies on the independence of participants, an unrealistic assumption common to many ML methods, despite it being well understood that there may be considerable cryptic relatedness in population cohorts; see, *e.g*., Fig. 3b in (***Bycroft et al., 2018***). A priori, it is unknown whether and how such structure affects effect size and interaction estimates. This casts doubt on the appropriateness of ML methods ignoring dependence, and on the reliance of parametric methods such as LMMs on restrictive assumptions in order to model dependence. Furthermore, deep learning methods, such as (***McCaw et al., 2022***), (i) require extensive hyper-parameter tuning that may be phenotype-genotype dependent, (ii) cannot take into account dependence structure amongst individuals, *i.e*., require independent and identically distributed (i.i.d.) training data, and (iii) have no mathematical guarantees to adequately control for type I errors or provide coverage at the desired levels.

Others have modelled non-linearities of allelic copies on trait based on additive-plus-dominance models and direct association tests, see *e.g*., (***Guindo-Martínez et al., 2021***). However, this approach still suffers from the aforementioned issues in the functional dependence of trait on population stratification and other covariates and confounders, as well as the potentially non-trivial dependence structure amongst the data of individuals.

Here, we introduce Targeted Genomic Estimation (TarGene), a method that accurately estimates effect sizes, pairwise and higher-order interactions amongst variants, and gene-environment interactions, by using flexible statistical and ML algorithms whilst equipping them with mathematical guarantees. TarGene does not suffer from any of the aforementioned shortcomings. Specifically, TarGene increases power for true discoveries, whilst simultaneously controlling false discoveries by: (i) Ensuring optimal bias-variance trade-off on the target quantity of interest (e.g. effect sizes or interactions), rather than merely optimising the fit of trait as a function of variants and covariates, and (ii) taking account of non-linear covariate effects, and (iii) non-linear genetic effects (additive/dominance). To our knowledge, other methods in the literature lack at least one or two of the above. For example, the standard additive-plus-dominance models (typically linear or LMM) can take into account the third aspect (iii) above, yet they do not take account of (ii) or offer guarantees for (i). We note that the gain in power, via (ii) using more advanced ML techniques (e.g. XGBoost, also employed by TarGene), over linear fits has already been demonstrated in (***McCaw et al., 2022***) on both simulations and UKBB data. That said, this paper explicitly mentions a short-coming of DeepNull/XGBoost: it cannot account properly for sample relatedness, which TarGene can. Furthermore, there are mathematical guarantees of type (i) for neither LMMs nor methods employed in (***McCaw et al., 2022***). So, in addition to improving power, TarGene both (a) incorporates relatedness and (b) guarantees reduction in residual model misspecification bias.

Fig. 2 summarises TarGene vs LMM-based approaches currently considered as gold standard in the population genetics literature for addressing the aforementioned complexities.

**Figure 2.**
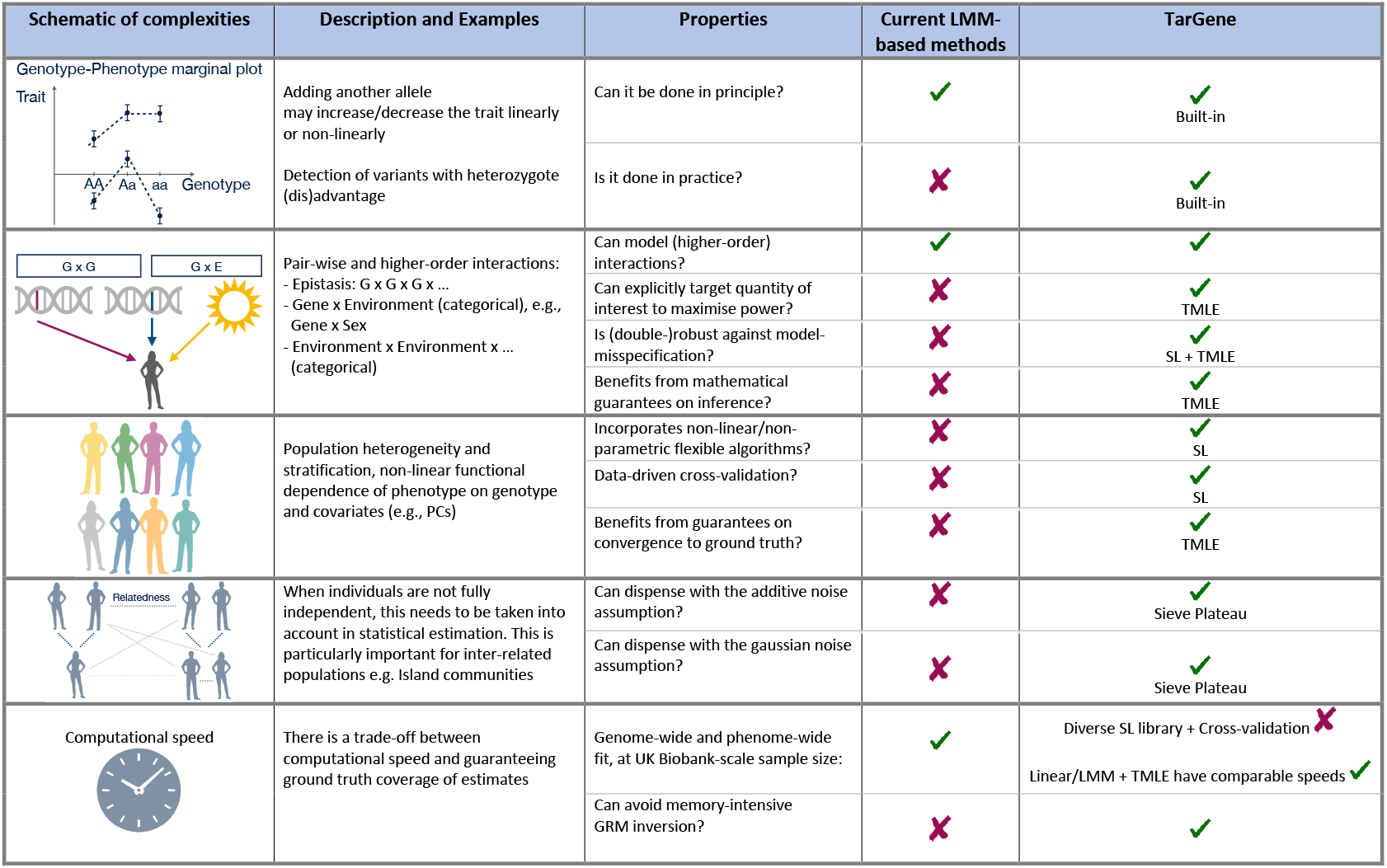
Comparison of TarGene vs current LMM-based methods considered as gold-standard in population genetics analysis. LMM-based methods use approximations that could result in reduced accuracy. In contrast, TarGene provides extensive flexibility and mathematical guarantees of ground truth coverage.

In brief, TarGene employs Super Learning, an ensemble machine learning method that estimates DNA variants’ effect sizes and strengths of interactions. It does so by data-adaptively (via *k*-fold cross-validation) combining a library of parametric and non-parametric methods (***van der Laan et al., 2007***), with the latter making no assumptions regarding the form of the phenotype-genotype relationship (***Benkeser and van der Laan, 2016***). The Super Learner (SL) combines methods to obtain a guaranteed optimal fit of the genotype-phenotype relationship. Any current or future, perhaps more powerful, estimation methods can be appended to its library of algorithms to improve performance. TarGene thus subsumes and supersedes any current model or ML based methods by incorporating them within the SL. Estimates are further enhanced by Targeted Maximum Likelihood Estimation (TMLE) (***van der Laan, 2017***). This step is crucial because it has the important qualities of being mathematically guaranteed to (a) reduce any residual bias due to model-misspecification, whilst (b) optimising the bias-variance trade-off, and (c) guaranteeing 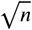-convergence as the sample size *n* increases (Fig. 1B). Finally, TarGene updates variance estimates via a network approach (***Davies and van der Laan, 2014***) to accurately account for population dependence structure, thus resulting in realistic p-values.

As a proof of concept, we apply TarGene to estimate the effect size of the *FTO* intronic variant rs1421085, a candidate causal variant for obesity (***Claussnitzer et al., 2015***) on all 660 binary and 118 continuous traits in the UK Biobank (UKB). We demonstrate a non-linear effect size, *i.e*., significantly differing effect sizes between the addition of the first and the second C allele, in 54 traits. This shows that TarGene can reveal non-linearity in genomic effect sizes whilst accounting for potentially complex heterogeneous population structures, as supported by its mathematical underpinnings.

To show the ability of the method to detect interacting variants, we consider the epistatic effect of six pairs of variants which were previously identified to be associated with hair color (***Morgan et al., 2018***). We find that three of those pairs are also epistatically associated with skin color, a correlated trait.

To demonstrate further applications of TarGene, we investigate variants potentially interacting via the vitamin D receptor (VDR). VDR is a nuclear hormone receptor that binds to calcitriol, the active form of vitamin D, and then forms a complex with the retinoid-X receptor (RXRA). We consider three genetic variants associated with differential expression of each of these three molecules. Thus, we estimate pair-wise and 3-point epistatic interactions amongst rs7971418, rs1045570, and rs3755967, affecting 660 binary and 118 continuous traits in the UKB. In the case of these variants, after multiple hypothesis testing, we do not find evidence of significant interactions. This is not unexpected because, evidently, detection of epistasis is extremely challenging (***Wei et al., 2014***).

## Results

We introduce TarGene in five steps, first highlighting shortcomings in current methods as we proceed using simple, easily interpretable simulations before explaining how TarGene addresses these problems. The mathematical guarantees behind TarGene are presented in the Methods section. As a proof of concept, we apply TarGene in two confirmatory settings and one exploratory study. For our confirmatory analyses, we estimate (i) the effect size of an *FTO* intronic variant, rs1421085, a candidate causal variant for obesity (***Claussnitzer et al., 2015***), as well as (ii) six epistatic effects, identified in (***Morgan et al., 2018***), on skin color. This exploratory study is motivated by the vitamin D receptor (VDR). Three variants (rs7971418, rs1045570, and rs3755967) were identified from their correlation with the abundance of molecules contained with this molecular complex. We investigate the pairwise interactions amongst those three variants and their 3-point interaction. In total, 660 binary and 118 continuous traits from the UK Biobank are investigated.

### Shortcomings in current methods

We illustrate two shortcomings of current approaches to GWAS, as well as their deleterious ramifications, via simulations and examples: (i) model-misspecification, and (ii) non-principled choice of confounders and covariate relations.

#### Model-misspecification in GWAS

In genome-wide association studies (GWAS), the effect size of a variant *V* on trait *Y* in the presence of confounders (such as sources of population stratification), is often referred to as the ‘beta’ coefficient. The following parametric form, or its LMM equivalent, is assumed to govern the data:

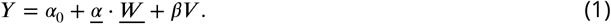

The effect size is then assumed to be equal to the coefficient *β*. However, in general this is not true because the data may be governed by a completely different probability distribution in which the effect size of the variant on trait is not equivalent to *β* above. The equation 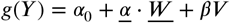 with link function *g*(*Y*) = *Y* and effect size *β* is used for a continuous trait *Y*. However, the same arguments and simulations apply to the logit link function *g*(*Y*) = logit(*Y*) for a binary trait *Y* in a case/control setting with *β* the log odds ratio of case to control.

To illustrate this important point, suppose a researcher assumes that the data follow Eq. 1 with 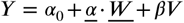. By this assumption, they thus declare the effect size of variant *V* on trait *Y* to be equal to *β* exactly. Nevertheless, they are only correct if the ground truth follows Eq. 1. If, instead, the ground truth differs, for example by addition of an interaction term between variant *V* and a confounder *W*_1_ (*i.e*., 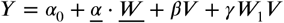) then the effect size of V on Y is made ambiguous: is the effect size *β* alone, or *γ*, or 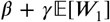? In a second example, if the ground truth has an exponential functional form *Y* = exp (*V* + (*V* + *ϵ*) *W*) (where *ϵ* = 0.2 and 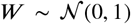 is assumed standard normally distributed) then, again, the true effect size is unclear. The answer is far from *β* = 1: it equals 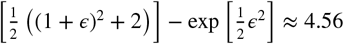.

We next used simulations to exemplify two ways in which model-misspecification, *i.e*., fitting data with a model different from the one from which it was generated, results in biased and statistically incorrect estimates. In Fig. 4, we present results from these simulations describing two distinct phenotype-genotype relations in the presence of confounders such as population stratification. To demonstrate the ubiquity of incorrect conclusions produced by incorrectly specified models, we independently varied over parameter space (1) the true effect size, (2) the noise on the data, and (3) the sample size, where the latter ranges from 1000 participants to a biobank-scale of 500, 000. These simulations show four features of model-misspecification (see Methods): (i) if the fitted model is far from the true data-generating distribution, then the slightest degree of measured correlation between *V* and *W* results in invalid inference of the effect sizes, even at smaller sample sizes; (ii) at any fixed level of noise, there always exists a sample size above which model-misspecification leads to invalid inference; (iii) replication is a necessary but not sufficient condition for declaring valid inference because fitting separate data samples drawn from the same distribution with the same (or a similar) misspecified model twice, results in equivalent invalid inference twice; and (iv) multiple hypothesis correction methods account for the testing of multiple hypotheses only, not for false discovery in a single hypothesis, such as those due to mis-specified parametric models. We conclude that it is crucial to avoid subjective modelling choices: model-misspecification is likely to give rise to invalid inference, especially when working with data sets as large as population biobanks which include complex population structures as well as gene-environment interactions (***Abdellaoui et al., 2022***).

#### Non-principled choice of confounders and covariate relations

Variables which confound the relationship between a variant *V* and a trait or disease *Y*, such as population stratification, deserve careful consideration. Usually, population genetics studies choose these variables subjectively. The extent by which such choices affect effect size estimation and hinder replication remains unclear. The UK Biobank project reports that inclusion of 16-20 principal components (PCs), labelled by self-reported ethnicity, is sufficient to capture sources of population stratification (***UKB, 2015***) (see also (***Canela-Xandri et al., 2018***; ***Neale, 2018***)). However, others (***Bycroft et al., 2018***) report use of up to 40 PCs and demonstrate significant population stratification across the entire UKB cohort. Consequently, for any particular analysis it is unclear what PCs to include, and whether by conditioning on PCAs (*e.g*., by including them as variables in GWAS fits) the true genetic signal is faithfully revealed. Additionally, there is no consensus on what covariate combinations (*e.g*., array batch or UKB assessment centre or sex × age^2^; see, for example, (***Canela-Xandri et al., 2018***; ***Neale, 2018***)) should be used.

We also discuss the additional issue of residual fitting sometimes employed in population genetics literature in Appendix Appendix B: Residuals of one fit as dependent variable in another fit. Residual fitting is the two-step process whereby the trait *Y* is first fitted linearly, either using linear regression or an LMM model, as a function of the selected covariates and random genetic effects. The residual of the fit, *i.e*., *Y* − *Y*_predict_, is then used a dependent variable in a second linear regression model, to estimate the effect size of the variants of interest. However, this procedure results in biased effect estimates and deflation or inflation of p-values (***Chen et al., 2018***).

### TarGene provides mathematical guarantees and realistic p-values

We next introduce our workflow, TarGene, and show that it does not suffer from any of the above shortcomings. We also illustrate the mathematical guarantees of coverage, asymptotically normal distribution, and realistic p-values our method provides.

TarGene is based on Targeted Learning (TL), a model-independent framework of estimation integrating causal inference, machine learning, and mathematical statistics and produces powerful estimators that are provably unbiased and efficient (***van der Laan and Rose, 2011***, ***2018***). TL consists of three steps: (1) Defining the quantity of interest model-independently; (2) Employing a diverse library of learning algorithms to learn the relevant portion of the true probability distribution, which results in an initial estimate of the quantity of interest; and, (3) Applying Targeted Maximum Likelihood Estimation (TMLE) to update and target the initial fit towards the quantity of interest, thereby reducing residual model-misspecification bias by optimising the bias-variance trade-off. Here, we briefly explain each of these steps (for more, see Methods) and show results of TarGene applied to UK Biobank data. In later sections we explain how TarGene accounts for (4) cohort population dependence, and (5) multiple hypothesis testing.

#### Step 1: TarGene defines the quantity of interest model-independently

In GWAS, whether the ground truth probability distribution *P*_0_(*y*, *v*, *w*) is linear, logistic, or has a more complicated form, the effect size of *V* on *Y* can be evaluated under any distribution *P*. By taking advantage of such model-independent definitions of both effect size and epistasis, TarGene resolves any ambiguity in estimation due to model-misspecification. Thus TarGene does not have to resort to any particular parametric assumptions. More specifically, effect size is otherwise known as the Average Treatment Effect (ATE) (***van der Laan and Rose, 2011***, ***2018***), and defined as

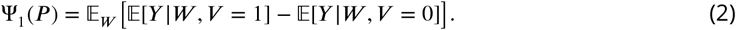

This is interpreted as the expected phenotypic change in *Y* for variant *V* = 1 relative to *V* = 0, whilst correcting for confounders *W* ; see also Fig. 1C.

It is common for genetic studies to examine additive models only. Nevertheless, because epistasis is crucial to understanding complex disease (***Mackay and Moore, 2014***), we similarly provide the model-independent definition of epistatic interaction between two variants *V*_1_ and *V*_2_ leading to variation in a trait or disease risk *Y* ,

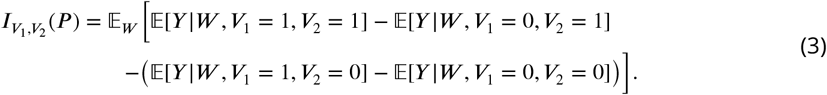

This definition was first introduced in (***Beentjes and Khamseh, 2020***); see also Fig. 1D. It is interpreted as the change in effect size of variant *V*_1_ on trait *Y* as variant *V*_2_ changes from 0 to 1.

#### Step 2A: TarGene data-adaptively incorporates confounders

Any population genetics analysis requires accounting for sources of population stratification in the specific cohort of interest. In the current literature, this is typically done by incorporating a number of Principal Components (PC) in a trait-independent analysis. Although standard, it is important that this analysis be performed for each database and repeated if a subset of the population in the cohort is used in order to obtain a lower bound on the number of PCs required. As a first example, in (***Bycroft et al., 2018***) the first 20 PCs were selected to account for population stratification based on labelling PC plots by self-reported ethnicity and visually inspecting their symmetry. This number may well be different for other more diverse cohorts, such as the *All of Us* (***Investigators, 2019***) cohort or the *Million Veterans Program* (***Gaziano et al., 2016***). As a second example, the analysis of the population structure in the whole UK Biobank cohort (***Sudlow et al., 2015***) may require far more PCs to account for population heterogeneity than the subset of individuals of White European ancestry as used in the current work (see Fig. 5, panels A-D).

To adopt a more principled approach to choosing confounding variables, TarGene incorporates data-driven methodologies that capture sources of population stratification confounding the relationship between *V* and *Y* (Fig. 1B). Specifically we select, in a data-adaptive manner, the optimal number of PCs for a given trait and set of variants (see Methods). In Fig. 5, panels A-D, we present a trait-independent PC analysis to construct a lower bound on the number of PCs required. Refining this to a trait-dependent analysis demonstrates that each trait has its own dependency on PCs, Fig. 5, panels E and F, and we recommend performing a sensitivity analysis by including higher-order PCs as part of the Super Learner (see below). Notably, if however supernumerary PCs do not strongly confound the variant-trait relationship, then the SL sets their coefficient to zero, thereby retaining only relevant PCs even if these are not consecutive in number. In Fig. 5C, we observe that the variant rs1421085 is randomised in the cohort and so population stratification captured by PCs is not a confounder of the phenotype-genotype relationship. This exercise is repeated for higher-order PCs and the other three variants considered in this manuscript (see Figs. 8–11). In this scenario, the only potential source of non-linearity in the functional dependence of phenotype on genotype and confounders is in the allelic copy (for a proof, see *Step 2A’* in Methods). Thus, a linear fit from *V* = 0 to *V* = 1 followed by a linear fit from *V* = 1 to *V* = 2 may yield equivalent effect sizes to TMLE up to the inference of confidence intervals. However, it is impractical to visually inspect the randomisation of each and every SNP on each and every trait in a typical statistical genetics analysis. For example, Fig. 5A and B show evidence of population stratification in the genotyped SNPs used to generate the PC plots. TarGene obviates the need for such manual inspections, irrespective of whether the variant is randomised or stratified in a cohort.

#### Step 2B: TarGene leverages a diverse combination of algorithms via Super Learning

It is unnecessary to expend computational resources on estimating the full probability distribution *P* (*y*, *v*, *w*) in order to evaluate the target parameters of Eqs. 2 and 3. Rather, only the parts 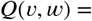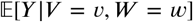 and *g*(*υ*, *w*) = *p*(*V* = *υ*|*W* = *w*) are required. TarGene leverages this by using a Super Learner (***van der Laan et al., 2007***), a stacking technique whereby various non-linear and/or non-parametric methods, neural networks, and tree-based algorithms in addition to parametric linear and logistic models more usually employed in GWAS, can be combined in a *k*-fold cross-validation scheme to find the optimal (in terms of mean squared error) combination of models to fit the data. Using *k*-fold cross-validation, SL is mathematically guaranteed to yield the combination of models with the best predictive power, as proven in (***van der Laan et al., 2007***). This procedure is depicted in Fig. 3, second panel.

**Figure 3.**
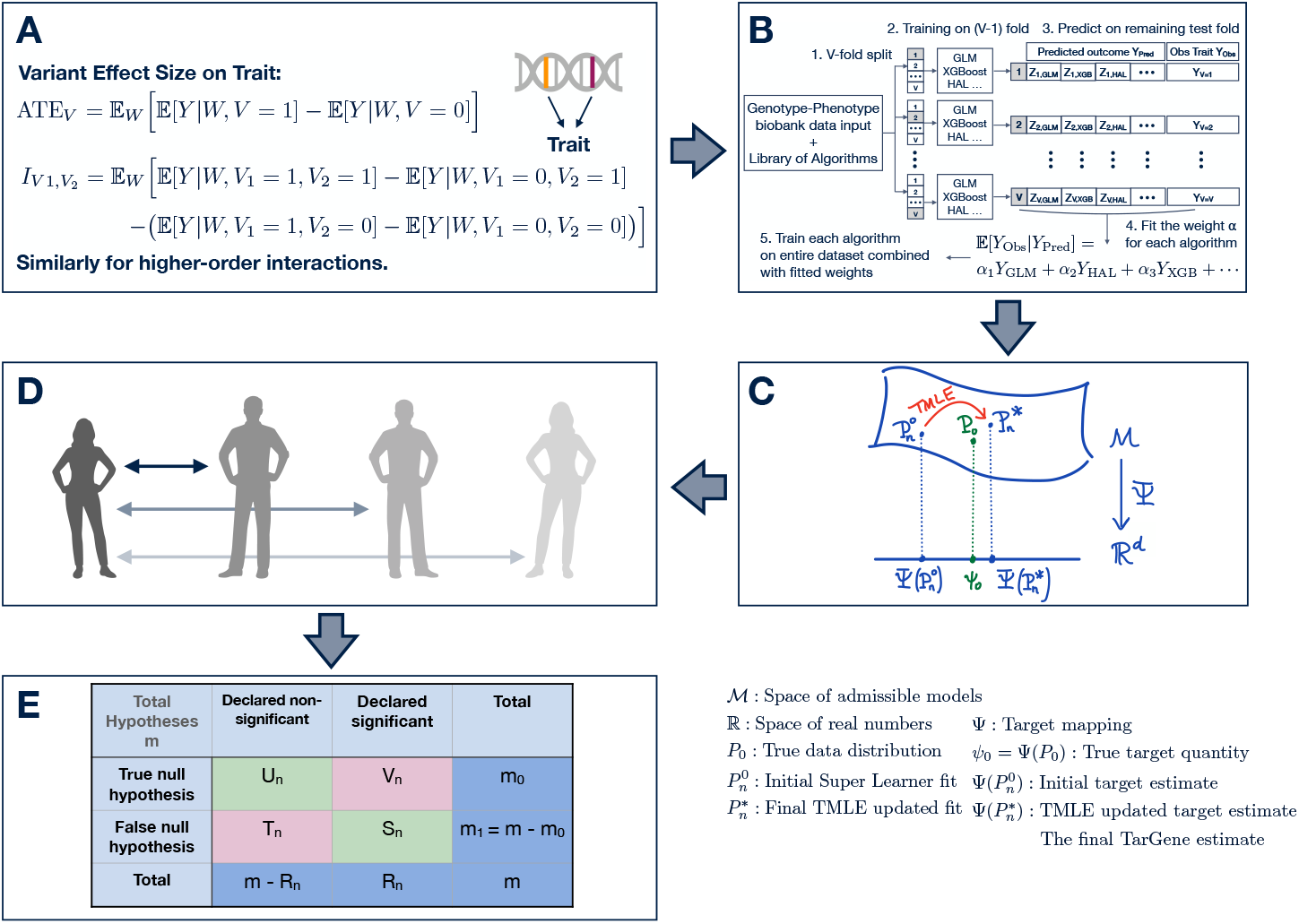
Workflow of TarGene. **A**: Model-independent definition of variant *V* effect size and two-point variant *V*_1_, *V*_2_ interaction effect on trait *Y*, taking into account sources of population stratification *W*. **B**: The Super Learner fit with a library of algorithms including linear, logistic, XGBoost and Highly Adaptive Lasso (HAL) algorithms. The SL is used to obtain an initial fit of the trait as a function of variant(s) and sources of population stratification. The user is free to add any other algorithm to this. *k*-fold cross-validation is performed to determine the algorithm or the combinations of algorithms with the lowest loss. **C**: Targeted Maximum Likelihood Estimation (TMLE) removes any residual model-misspecification bias for the target quantity of interest, resulting in optimal bias-variance trade-off with mathematical guarantees. **D**: Uncertainties in step C are updated to take into account dependences among individuals in the population. **E**: Multiple hypothesis testing is performed.

In general, the output of SL is an initial estimate 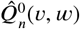 of the function *Q*(*υ*, *w*), as well as an initial estimate of the target parameter obtained by plugging 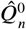 into Eq. 2:

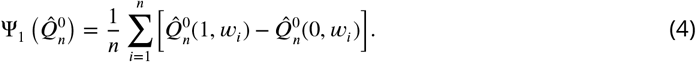

The average is taken over the cohort of size *n*, and *w*_*i*_ are the covariates of participant *i*.

#### Step 3: TarGene performs a targeted update via TMLE to optimise bias-variance

SL is optimised to produce the best estimate, 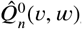, of the function 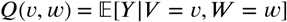. However, it is not optimised for estimating the DNA variant’s true effect size on phenotype, *i.e*., the target parameter Ψ_1_(*P*_0_). As a result, there may be residual bias in the initial estimate Eq. 4, *i.e*., a discrepancy between the initial effect size estimate, 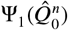, and its true value, Ψ_1_(*P*_0_). Under mild assumptions, mathematical theory (***van der Laan, 2017***) allows us to separate this discrepancy into three components (see Eq. 24 in Methods). The first component represents residual bias due to model-misspecification. The TMLE update is mathematically guaranteed to reduce this bias component in order to achieve optimal bias-variance trade-off on the final target quantity of interest.

The second component represents the variance on the estimate which is used to provide final 95% confidence intervals. The third, and final, component arises from finite sample size and is guaranteed to shrink at rate 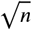 as the sample size *n* increases provided the SL is rich enough. Moreover, together with the vanishing of the first component, TMLE makes the final estimate asymptotically normally distributed.

To perform the targeted update step of the TL framework, the TMLE step, requires an estimate of the *propensity score* or *treatment mechanism g*(*υ*, *w*) = *p*(*V* = *υ*|*W* = *w*). In the context of GWAS, this is the probability that an individual carries variant *v* given that they belong to population stratum *w*. In particular, if *g*(*υ*, *w*) is essentially independent of population stratum, then variant *V* is *not* stratified in the population. To verify this, the test for the null hypothesis *H*_0_ ∶ *g*(*υ*, *w*) = *p*(*V* = *υ*) is easily incorporated into the TL framework. In practice, the propensity score is also estimated via Super Learning as discussed in the previous section.

Since the UK Biobank contains over 450, 000 samples, it is likely that the asymptotic regime is reached in this data set. To demonstrate the behaviour of TMLE as compared to biased estimation methods, we performed a bootstrap analysis on the UK Biobank (Fig. 6). In this precise example, we estimated the effect of substituting TT with TC for rs1421085 on BMI. From the plot, it is notable that the effect size distributions provided by the bootstrap SL (orange) and the linear-model (grey) do not overlap. This is a real-data example of the model-misspecification phenomenon illustrated in the schematic in Fig. 1 A and B. Adding an extra targeting step in addition to any of these methods, reduces the residual bias and always brings estimates back to the same confidence region (see Figure legend for details).

#### Step 4: TarGene accounts for population dependence structure

Above we describe how to estimate the effect size of one DNA variant on a single phenotype measured on independent and identically distributed data. However, if the data are dependent, for example because of genetic relatedness (***Naseri et al., 2021***; ***Powell et al., 2010***), care must be taken when estimating the variance on the estimates. In current practice, the genetic dependence between individuals is accounted for by incorporating the genetic relationship matrix **G** (GRM) as random effects in a linear mixed model (LMM):

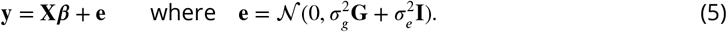

Here 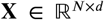 is the matrix of fixed effects, *d* the number of fixed effects, and ***β*** the fixed effect sizes. There are two random components: (i) 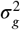 denoting the magnitude of the genetic variance, and (ii) 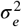 denoting the magnitude of the residual variance. There is little justification for this model’s strong restrictions. Besides model-misspecification, adopting an LMM may be invalid for three reasons: (i) The error term **e** need not be normally distributed (***Fusi et al., 2014***); (ii) The variance of **e**, 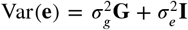, need not decompose additively into one part due to population stratification and another due to residual environmental, technical, and other noise; and, (iii) The complexity of population stratification need not be captured by a single parameter 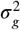 multiplying the GRM.

In TarGene, we neither assume individuals are independent nor impose the strong restrictive assumptions of an LMM mentioned above. Instead, we adopt a network approach, drawn from (***Davies and van der Laan, 2014***), to incorporate the genetic dependence of individuals model-independently by taking into account ancestral diversity and familial relatedness amongst individuals as reported, *e.g*., in Fig. 3b of (***Bycroft et al., 2018***). The method calculates the variance of the effect size target parameter by constructing Sieve Plateau (SP) variance estimators (see Methods) that incorporate genetic dependence among individuals.

In brief, the SP estimator computes a variance estimate for a range of thresholds *τ*, by considering individuals to be genetically *independent* if their genetic distance exceeds *τ*. The genetic distance between a pair of individuals (*i*, *j*) equals 1 − GRM_*i*,*j*_, *i.e*., one minus their genetic related-ness value. As the distance threshold *τ* increases, fewer individuals are assumed to be genetically independent. For instance, the estimate corresponding to a distance of *τ* = 0 corresponds to the i.i.d. hypothesis, while a distance of *τ* = 1 incorporates pairs of individuals who are not genetically correlated. TarGene varies the threshold *τ* from 0 to 1 and fits a curve to the corresponding variance estimates. The maximum of this curve is the most conservative estimate of the variance of the target parameter estimator and constitutes our corrected variance estimator. In Figure 6, we show the effect of this correction for the effect sizes obtained for rs1421085 on all parameters under investigation (see Application of TarGene to an FTO variant). Since, the correction can only increase the variance estimate, we only correct those for which the associated p-value is under our decision threshold (0.05). The p-values resulting from both the i.i.d. (red) and the sieve variance plateau estimators (blue) are reported. For 19 parameters (≈ 5%), the corrected estimate changes the hypothesis test decision at the 0.05 threshold.

#### Step 5: TarGene controls for multiple hypothesis testing

When testing multiple hypotheses simultaneously in order to answer a question of interest, it is essential to state explicitly which error rate of false positives one seeks to bound, and then choose the multiple hypothesis correction procedure that affords maximal power (*i.e*., fewest Type II errors) whilst bounding the desired Type I error rate. Since TarGene produces asymptotically normal estimators described as empirical means of efficient influence functions, the theory is sufficiently rich to be combined with any desired definition of false positives, *e.g*., the family-wise error rate (FWER) or the false discovery rate (FDR) (see Methods). In this work, we use the marginal step-down Benjamini–Hochberg procedure of (***Benjamini and Hochberg, 1995***) to control the FDR at ≤ 0.05. However, researchers can combine TarGene with any multiple testing procedures to control FWER, FDR, or other error rates, for marginal or joint multiple hypothesis testing, as described for example in (***Dudoit and van der Laan, 2008***).

### Application to the UK Biobank

To investigate the benefits of TarGene, we performed three distinct analyses that are detailed below. The first aims at contrasting our approach with the gold standard LMM’s method on a well studied variant. The second, displays the ability of TarGene to replicate pairwise interactions previously reported for hair colour (***Morgan et al., 2018***). The third, more exploratory, investigates whether epistatic relationships exist amongst multiple loci related to vitamin D receptor (VDR) biology. We present here our major findings after the targeting step, sieve variance correction and multiple hypothesis adjustment. Apart from the second analysis, we consider the 776 UK Biobank traits defined by GeneATLAS (***Canela-Xandri et al., 2018***).

#### Application of TarGene to an FTO variant

In order to demonstrate our method we performed a phenome-wide association study (PheWAS) using UK Biobank. We chose a well studied variant, rs1421085, located in the first intron of the *FTO* gene. The T to C nucleotide substitution has been predicted to disrupt the repression of *IRX3* and/or *IRX5*, thereby leading to a developmental shift from browning to whitening programs and loss of mitochondrial thermogenesis (***Claussnitzer et al., 2015***). This variant has also been associated with several related traits such as BMI and obesity (***Frayling et al., 2007***). To allow TarGene to discover non-linear effect sizes as previously reported in (***Wood et al., 2016***), the analysis was not restricted to the substitution of one T allele to C. Instead, we investigated all three changes, namely TT → TC, TC → CC and TT → CC. We adjusted p-values to control for the false discovery rate using the Benjamini-Hochberg method. A summary of all estimation results is provided in Supplementary Table 1. Each change requires a dedicated targeted estimate as each corresponds to one of three different target quantities of interest. This is in contrast with a linear model assuming that these quantities are equal to *β*, *β*, and 2*β* respectively.

Of 776 traits under investigation, 20.7% are reported as significantly associated to rs1421085 by GeneATLAS. Using TarGene, we find the following for the three quantities of interest (FDR ≤ 0.05, Benjamini–Hochberg):

- TT → TC *only*: 9.1% of traits are significant.
- TT → TC *or* TC → CC: 12.5% traits are significant.
- TT → TC, TC →CC *or* TT →CC: 15.4% traits are significant.

The distribution of p-values obtained via TarGene is shifted towards less significant values as compared to the GeneATLAS analysis (see Fig. 6 in Supplementary figures). TarGene finds fewer significant results than GeneATLAS but provides mathematical guarantees of statistical coverage of ground truth on the results it finds, leading to fewer false positives whilst maximising power.

Notably, because we investigate each allelic change as a separate quantity, TarGene can also find associations that are undetectable by linear models such as LMMs. In Eq. 2 we define the effect size of a SNP *V* on a phenotype *Y* model-independently via the target parameter Ψ_1_(*P*). This is the effect on phenotype when a single allelic copy is present (*V* = 1) versus when there is none (*V* = 0), *e.g*., TC vs TT. However, there is another equally valid, and potentially distinct, way of describing the effect of an additional allelic copy on *Y*, namely the effect on phenotype when two allelic copies are present (*V* = 2) versus one (*V* = 1), *e.g*., CC vs TC. The corresponding model-independent definition is

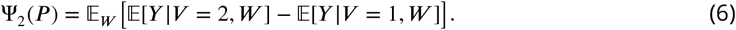

Conventional GWAS, such as LMMs (***Lippert et al., 2011***), assume linearity of genetic effect on phenotype (*i.e*., assume Ψ_1_(*P*) = Ψ_2_(*P*)). However, there are no compelling biological or mathematical reasons why the two effect sizes should be equal (***Veitia, 2003***). More recent methods, such as KnockoffGWAS (***Sesia et al., 2021***), do not estimate effect size at all.

TarGene avoids this unnecessary assumption by estimating both effect sizes separately. This does not necessarily increase the burden of multiple hypothesis testing because one can choose to query the effect size Ψ_1_(*P*) at the TMLE step for each trait-variant pair. Alternatively, if one specifically wishes to identify non-linear effect sizes of trait-variant pairs, or classify the type of DNA variants and/or phenotypes for which such non-linearities occur, one can directly estimate the difference Ψ_2_(*P*) − Ψ_1_(*P*) (Methods).

The two scenarios under consideration are (i) the change TT → TC, and (ii) the change TC → CC. We find 54 traits for which rs1421085 displays significant non-linear effect sizes, 40 of which are highly correlated with BMI. For instance, we find that the departure from homozygous TT to heterozygous TC is associated with an increase of 0.77 kg (95% CI: 0.68 − 0.85). In comparison, the departure from heterozygous TC to homozygous CC is associated with a significantly larger increase of 1.31 kg (95% CI: 1.19 − 1.43). For illustration, a subset of significant non-linear traits is presented in Fig. 6; Supplementary Table 2 contains the complete list. As might be expected, estimates reported by GeneATLAS, based on a linear LMM approach, fall in-between estimates from our two scenarios, representative of an averaging effect.

Notably, for urticaria and erythema, replacing a single T with C (*i.e*., TT → TC) is associated with significantly (*p* = 2.9 × 10^−4^) *lowered* risk whereas replacing the second T with C (*i.e*., TC → CC) is associated with significantly (*p* = 9.5 × 10^−3^) *elevated* risk (Fig. 6, middle). Thus TarGene can capture variant-trait pairs displaying *Heterozygote Advantage* (***Hedrick, 2012***). Such patterns cannot be detected by a linear model assuming equal allelic effect sizes.

These two results have shown that model misspecification and the curse of dimensionality (see Fig. 1) can be problematic in two ways: (i) Significant results may be reported that are false positives and may thus result in wasted experimental follow-up studies, and (ii) significant hits may not be reported that might have otherwise informed future research, *i.e*., they are in fact false negatives. Note that linear models are a standard part of the SL library of TarGene so if statistical inference using a linear model is correct, then TarGene will choose the model in a data-driven manner. TarGene provides mathematical guarantees of coverage of ground truth thus, in combination with multiple hypothesis testing, bounding false positives whilst minimising false negatives.

#### Replication of Epistatic effect sizes

Detection of epistatis in complex traits can be challenging (***Wei et al., 2014***). For example, epistatic interactions are expected to be much smaller than main effect sizes, which can already be small for polygenic traits. It is therefore crucial that an optimal bias-variance trade-off is employed when estimating epistatic interactions. We employ TarGene to estimate epistasis in a model-independent way, see Eq. 3 and adjacent discussion. An interesting comparative ground for TarGene is found in (***Morgan et al., 2018***), a study investigating hair color, in which six pairs of SNPs were reported to be statistically interacting using a logistic regression model and a likelihood ratio test. We note however, that the likelihood ratio test statistic has no direct comparison with the interaction effect we report. Also, hair colour is not a trait investigated by GeneATLAS. Since hair and skin color have been shown to co-vary (***Sulem et al., 2007***), we focused instead on the latter which is part of GeneATLAS Out of the six reported significant pairwise interactions for hair color, we replicate three for skin color (see Supplementary Table 3), namely: rs1129038 and rs1805008 (p-value ≈ 2.43 × 10^−8^), rs117119427 and rs1805009 (p-value ≈ 4.57 × 10^−6^) and rs1805007 and rs6088372 (p-value ≈ 6.11×10^−7^). This demonstrates the ability of TarGene to detect epistatic interactions. The reason why not all six interactions are replicated could be a consequence of model misspecification of the logistic regression in (***Morgan et al., 2018***), but is likely also affected by the non-perfect relationship between hair and skin color.

#### Exploration of new epistatic loci

Next we investigated potentially interacting variants involved in VDR biology. VDR is a nuclear hormone receptor that binds to calcitriol, the active form of vitamin D, and forms a complex with the retinoid-X receptor (RXRA). This complex can then enter the nucleus and bind to specific genetic domains to regulate transcription of many genes. Because this mechanism depends on three interacting molecules (calcitriol, VDR and RXRA), it is a natural field of investigation for epistasis. We thus identified three genetic variants that have been associated with differential expression of each molecule in turn. From [eQTLGen](https://www.eqtlgen.org/), A to C change of rs7971418 is associated with increased levels of VDR; G to T change of rs1045570 is associated with increased levels of RXRA; and, C to T change of rs3755967 has been associated with a decreased level of calcitriol (***Jiang et al., 2018***). For each SNP, we restricted our attention to the departure from homozygosity of the major allele to heterozygosity and investigated all three pairwise interactions as well as the 3-point interaction. We provide estimation results and p-values for pairwise interactions in Supplementary Table 4, and for 3-point interactions in Supplementary Table 5. In each case, we also provide adjusted p-values to control the FDR (via the Benjamini-Hochberg method) across all traits being tested for each SNP combination. Although 43, 39 and 36 pairwise interactions (for rs1045570 and rs3755967, rs1045570 and rs7971418, rs3755967 and rs7971418 respectively), as well as 29 3-point interactions, were found significant prior to multiple testing correction, no interaction was significant following FDR correction at the 0.05 threshold. This is not unexpected because detection of epistasis is extremely challenging (***Wei et al., 2014***). However, this analysis demonstrates the opportunities that our method provides for the general investigation of higher order interactions.

## Discussion

We have introduced TarGene, a workflow for targeted estimation of variant effect sizes and epistatic interaction effects on polygenic traits, which dispenses with unnecessary assumptions currently widespread in the statistical genetics literature. TarGene consists of five steps: (1) Defining the quantity of interest, *e.g*., effect size or interaction, model-independently; (2) Employing a diverse library of learning algorithms, such as data-adaptive algorithms with proven convergence properties (***van der Laan, 2017***), to learn the relevant portion of the true probability distribution, which results in an initial estimate of the quantity of interest; and, (3) Applying Targeted Maximum Likelihood Estimation (TMLE) to update and target the initial fit towards the quantity of interest, thereby reducing any remaining bias to optimise the bias-variance trade-off on the quantity of interest. This is followed by (4) variance correction for cohort population dependence, and (5) multiple hypothesis testing. The estimators TarGene produces are context-independent, *i.e*., are applicable beyond variant-variant interaction, to any discrete set of variables affecting an outcome of interest, *e.g*., interactions of variant × sex, or any other binary or discrete environmental factors (***Abdellaoui et al., 2022***).

TarGene offers a number of distinct advantages over current commonly employed LMM approaches, as summarised in Fig. 2. In particular, since it is firmly rooted in the mathematical estimation framework of Targeted Learning, TarGene avoids model-misspecification bias, produces asymptotically normal and efficient estimates, and is doubly-robust. Furthermore, due to the flexibility of its SL libraries and the TMLE step, it can be readily applied to more ancestry heterogeneous biobanks such as *All of US* (***Investigators, 2019***) or the *Million Veterans Program* (***Gaziano et al., 2016***), as well as more strongly inter-related cohorts such as island communities. TarGene also allows for integration of data from multiple biobanks. We will pursue the application of TarGene to such meta-analyses in a future work.

The strength of TarGene lies in bespoke analyses of effect sizes and interactions amongst targeted variants of interest, providing mathematically guaranteed coverage of the ground truth. The run time of this workflow depends on how precise and unbiased a researcher wishes to be regarding the answer to their question of interest. For researchers interested in applying TarGene for genome-wide studies across multiple traits, we note that there is a trade-off between computational speed and guaranteeing ground truth coverage of estimates. In such a scenario we therefore recommend equipping the SL with computationally light algorithms only, such as a linear model, GLMnet or LMM, reducing the cross-validation burden significantly, before running the TMLE step. The lightest version of TarGene can be run in 10 hours on a HPC cluster, for a GWAS with 600,000 variants. See Table 2 for details. Significant hits from an initial light run can then be verified by a more comprehensive TarGene run with a stacked SL library. Taking advantage of the approach described in (***Tuglus and van der Laan, 2011***) nevertheless allows the researcher to control the final FDR of this two-stage procedure. We also recommend applying the SP variance correction to estimates with p-values near the FDR threshold to ensure their robustness. We also remark that, although the SP variance estimator requires reading the GRM into memory, in comparison to LMMs, TarGene does not require the memory-intensive inversion of the GRM. This inversion step may be prohibitive for UKB-scale cohort sizes, depending on institutional resources.

In future work, we plan to explore non-linearities in variant allelic copies on trait, which as of yet have not been systematically explored in the literature for either homogeneous or diverse populations. We will also investigate the contribution of epistatic interactions of specific variants on various polygenic traits for a variety of biological mechanisms.

## Methods

### TarGene provides mathematical guarantees and realistic p-values

We present the three main steps in the model-independent estimation framework of Targeted Learning (TL) (***van der Laan and Rose, 2011***, ***2018***) in detail: (1) Defining the quantity of interest model-independently; (2) Super Learning; and, (3) Targeted Maximum Likelihood Estimation (TMLE). In later sections we detail how TarGene (4) incorporates population dependence in the cohort, and (5) accounts for multiple hypothesis testing.

#### Step 1: TarGene defines the quantity of interest model-independently

The effect size of a DNA variant *V* on trait *Y*, correcting for population stratification via confounders *W*, is defined as

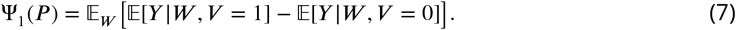

This is interpreted as the difference between the expected phenotype if the variant *V* = 1 versus *V* = 0, whilst correcting for confounders. The ground truth probability distribution is called ***P***_0_. The true (but unknown) effect size is denoted by *ψ*_1,0_ = Ψ_1_(*P*_0_).

The epistatic interaction between two variants in their effect on a given trait or disease *Y* has been defined in (***Beentjes and Khamseh, 2020***). This model-independent definition is

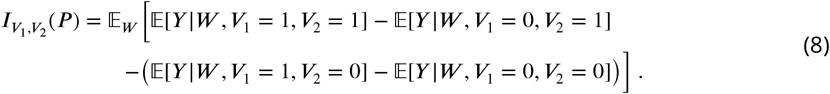

The genomic interpretation of this definition is: “Having correctly adjusted for confounders, is the effect of variant *V*_1_ on trait modulated by the status of variant *V*_2_ and, if so, by how much and with which sign?” The ground truth interaction is denoted 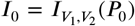. This definition of 2-point interaction, which is an extension of ATE with more than one variant, has been further generalised to higher-order interactions amongst *n* variants (***Beentjes and Khamseh, 2020***).

As an example, suppose the ground truth trait ***Y*** has the following expectation value:

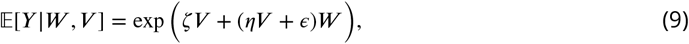

where *V* represents the variant of interest, W represents other covariates, *e.g*., principal components, and *ζ* and *η* represent different parts of the parameter space. Suppose also that *W* is distributed according to the normal distribution 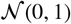. Since we are interested in measuring the effect size of *V* on trait *Y*, let us consider the terms appearing in the computation of the log-odds ratio or ATE effect size:

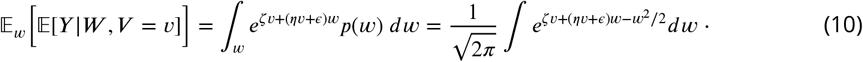

Completing the square and calculating the integral gives:

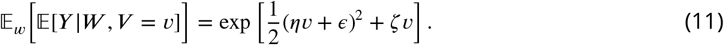

Therefore, the ground truth effect size is:

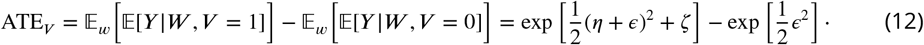

Now, suppose that we fit the ATE using the following misspecified model for expectation value, assuming linearity:

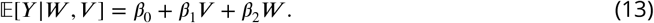

Then the ATE has the following expression in terms of the model parameters:

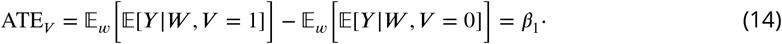

##### Simulation examples of model-misspecification

We generated data from the ground truth, in this case Eq. 9 with *W* standard normally distributed as stated, at a given value of *ϵ* = 0.3 without loss of generality. Here *ϵ* controls the levels of heteroskedasticity in the data as *V* varies from 0 to 1. We perform simulations in two scenarios:

1. The variant *V* and the source of population stratification *W* are independent
2. The variant *V* depends on *W* (*e.g*., principal components, location, batch, …). For simplicity, we consider the following dependence structure:

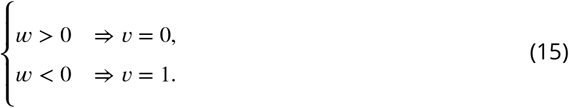

When 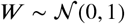, this leads to ≈ 80% correlation between *V* and *W*.

We illustrate the issue of model-misspecification using simulations based on parametric models that are then fitted (i) with the correct parametric form, and (ii) with a misspecified parametric model. We perform these simulations probing different parts of the parameter space and at various sample sizes.

###### Simulation 1 (top panels of Fig. 4)

We generate data from the ground truth distribution given by the exponential model explored in the previous section Eq. 9,

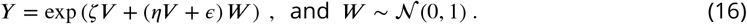

under three different scenarios: (a) the variant *V* is generated using a binomial distribution, and completely independent of *W*, (b) the variant *V* is generated by dichotomising *W* using an arbitrary cut-off, and (c) similar to (b) but also adding various degree of Gaussian noise when generating *V* from *W*. In scenarios (b) and (c) the variant *V* has various degrees of dependency on *W*, which is to be expected as *W* here represents sources of population stratification. Without loss of generality we set *ζ* = −1, and vary *η* and *ϵ* over a range of values to explore different part of the parameter space. More specifically, we probe the parameter space by setting *η* = [−2, 2] in steps of 0.5 and *ϵ* = [0, 1] in steps of 0.125. In each case, the ground truth exponential model is fitted with the misspecified linear model:

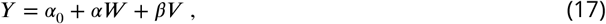

to obtain the assumed effect size *β*.

**Figure 4.**
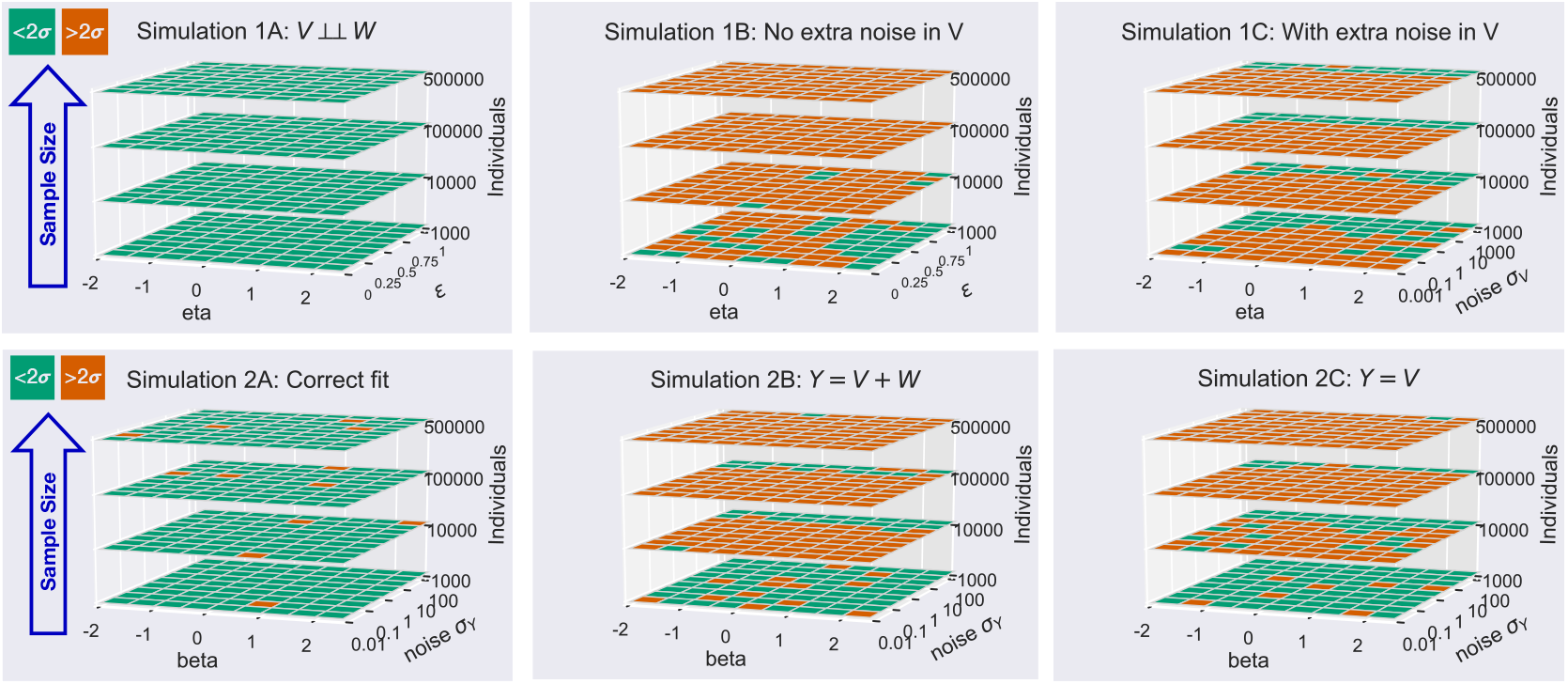
Model misspecification can yield biased estimates. In simulations, estimated effect sizes are either within (green) or outside (red) the 95% confidence interval. The striking conclusion of these simulations is that, when the model is misspecified, almost all estimates are invalid even at moderately large sample sizes. Top and bottom panels represent two distinct sets of simulations involving variables *V*, *W* and *Y* representing variant, sources of population stratification and trait, respectively. **Top panels**: Trait *Y* is generated from *V* and *W* via an exponential model and yet is fitted with a linear model (see Simulation examples of model-misspecification). Simulation **1A**(left): When *V* and *W* are *completely* independent (no population stratification present), the estimated effect size is correct (indicated in green) even when a misspecified linear model is fit to the exponential distribution. This result is irrespective of sample size. Simulation **1B**(middle): When, in a more realistic case where population stratification is present, *V* and *W* are dependent, then effect size estimates are incorrect (shown in red) when a misspecified linear fit is employed. This issue is exacerbated as the sample size grows, and manifests even with the slightest degree of Pearson or Spearman correlation (see Methods). Simulation **1C**(right): High levels of noise *σ*_*V*_ hide dependence between *V* and *W* so inference at small sample sizes may be within 1*σ*. As sample size grows, model-misspecification is increasingly exposed. **Bottom panels**: The trait *Y* is generated from *V* and *W* via a polynomial model yet fitted with a misspecified polynomial model (see Methods). In misspecified models, effect size estimates become more incorrect as sample size increases. In all subfigures the true data distribution is generated according to the polynomial model in Eq. 18. The plots indicate *β* estimated using three different models, taking values in different parts of the parameter space. Simulation **2A**(left) indicates the estimated effect size where the true model is used. When there is no model misspecification, estimates are valid irrespective of sample size. In Simulations **2B**(middle) and **2C**(right) two misspecified models are used to estimate *β*.

###### Simulation (1A)

In the unrealistic scenario where *V* and *W* are *truly* independent, a misspecified linear model happens to coincide with the true effect size for all values *ϵ* and *η* (Fig. 4, top panel, 1A). This is the case even though the goodness-of-fit measurements clearly indicate the non-linearity of the data, *e.g*., *R*^2^ ≈ 0.3 and extremely high-values of the Jarque-Bera index, indicating non-normality of the data. The previous statement holds irrespective of sample size. This behaviour has been observed in randomised control trials (RCTs) where the treatment mechanism is truly randomised with respect to known confounders and is therefore independent of these confounders (***Rosenblum and van der Laan, 2009***). This cannot be said for GWAS where sources of population stratification, *e.g*., genetic ancestry, are clearly visible from PCA projections in variant space.

###### Simulation (1B)

The variant *V* is generated by dichotomising *W* using an arbitrary cut-off to induce dependence between *W* and *V*. Without loss of generality, the cut-off is chosen such that *V* = 0 if *W* > −3.0 and *V* = 1 if *W* < −3.0. The conclusions below are similar irrespective of the choice of cut-off, which was tested to range from −3.5 to 3.0 in steps of 0.5, or larger values of *ϵ*. The traits are again generated using Eq. 16. As an example, for sample size 10, 000 (Fig. 4, top panel, 1B), the effect sizes estimated by the misspecified linear model are mostly incorrect. Furthermore, in about 50% of the cases, even the sign is inferred incorrectly, where a positive effect size of a variant on trait is estimated to be negative, instead of positive, when fitted with a misspecified model. We also note that the dependence structure in this example is not captured by either Pearson (≈ 0.1) or Spearman (≈ 0.1) correlations, both indicating a weak degree of correlation.

###### Simulation (1C)

This scenario is similar to (1B), following Eq. 16, but also adding various degrees of Gaussian noise when generating *V* from *W*. The noise is specified to take on values [0.001, 0.01, 0.1, 0.5, 1.0, 5.0, 10, 50, 1000]. As observed in Fig. 4 (top panel, 1C), at fixed sample size, large levels of noise may hide the dependency between *V* and *W*, resulting in effect sizes that match the ground truth value. However, as the sample size increases, the level of noise has to be extreme (50 or 1000) to cover for the misspecified model.

The simulations show that (i) at any fixed level of noise, there exists a sample size *N* for which model-misspecification leads to invalid inference due to shrinking variance and, therefore, (ii) when working with large data sets, such as the UKB, it is crucial to avoid subjective modelling choices as model-misspecification is likely to give rise to invalid inference.

###### Simulation 2 (bottom panels of Fig. 4)

There is nothing special about the choice of the exponential data generating distribution in Simulation 1. To exemplify this, we perform another set of simulations with data generated from a simple polynomial model:

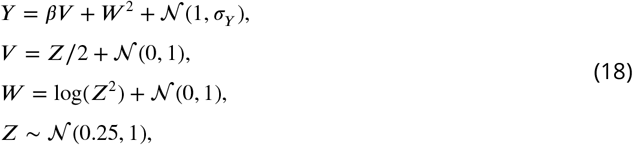

where *Z* is an auxiliary variable which is used to generate dependency between *V* and *W*. The independent noise in *Y*, *i.e*., *σ*_*Y*_, is varied by setting it to 0.01, 0.1, 1, 10 and 100. Without loss of generality, the true effect size *β* is probed from [−2, 2] in steps of 0.5. Fig. 4 (bottom panel, 2A), indicates the correct estimated effect sizes when the true model is used. As expected, most results are within two standard deviations, there are only a few values (≈ 4%) more than two standard deviations away from the ground truth.

###### Simulation (2B)

Here, the data are fitted with the following misspecified model:

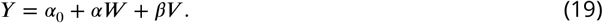

The results in Fig. 4 (bottom panel, 2B), clearly indicate that model-misspecification becomes more manifest as the sample size grows. At 10, 000 samples most estimated values are incorrect. At the UKB size, almost all estimated effect size values are incorrect.

###### Simulation (2C)

Here, the data are fitted with the following misspecified model:

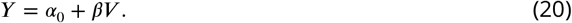

The results are presented in Fig. 4 (bottom panel, 2C), with the same conclusions as above.

We furthermore note that replication of results, even on two different databases, does not guarantee that estimated quantities and their confidence intervals span the ground truth, when a misspecified model is used. Therefore, under the assumptions that two data sets have a similar distribution, replication should be treated as a necessary but not sufficient condition for accuracy of the results: Fitting separate data samples drawn for the same distribution with the same (or similar) misspecified model twice, results in an equivalent invalid inference twice. Finally, we recall that multiple hypothesis correction methods correct for the testing of multiple hypotheses, not for false discovery in a single hypothesis, such as those predicted via misspecified parametric models.

#### Step 2A: Variant preprocessing and PC analysis

For the analysis in Fig. 5, we started with all directly genotyped variants in UKB and applied LD block removal, ± 10 Mb around rs1421085 position, using PLINK2 (***Chang et al., 2015***; ***Purcell and Chang, 2021b***), with *R*^2^ = 0.1. We filtered variants based on Minor Allele Frequency (MAF) threshold > 0.05 and performed LD pruning using PLINK (***Purcell et al., 2007***; ***Purcell and Chang, 2021a***) (1000kb window, 50 variants step size, and *R*^2^ = 0.05). We used the 33,483 remaining biallelic genotyped variants for 452,149 self-reported white individuals as input to FlashPCA2 to perform a partial PCA. Labelling the PCA plots by sex, age and batch, and plotting the corresponding cumulative distributions did not indicate any structure for these variables. A slight degree of separation by assessment centre was observed in PC1-2 cumulative distribution plots. Self-reported ethnicity is the dominant driver for the PCs as observed in Fig. 5.

**Figure 5.**
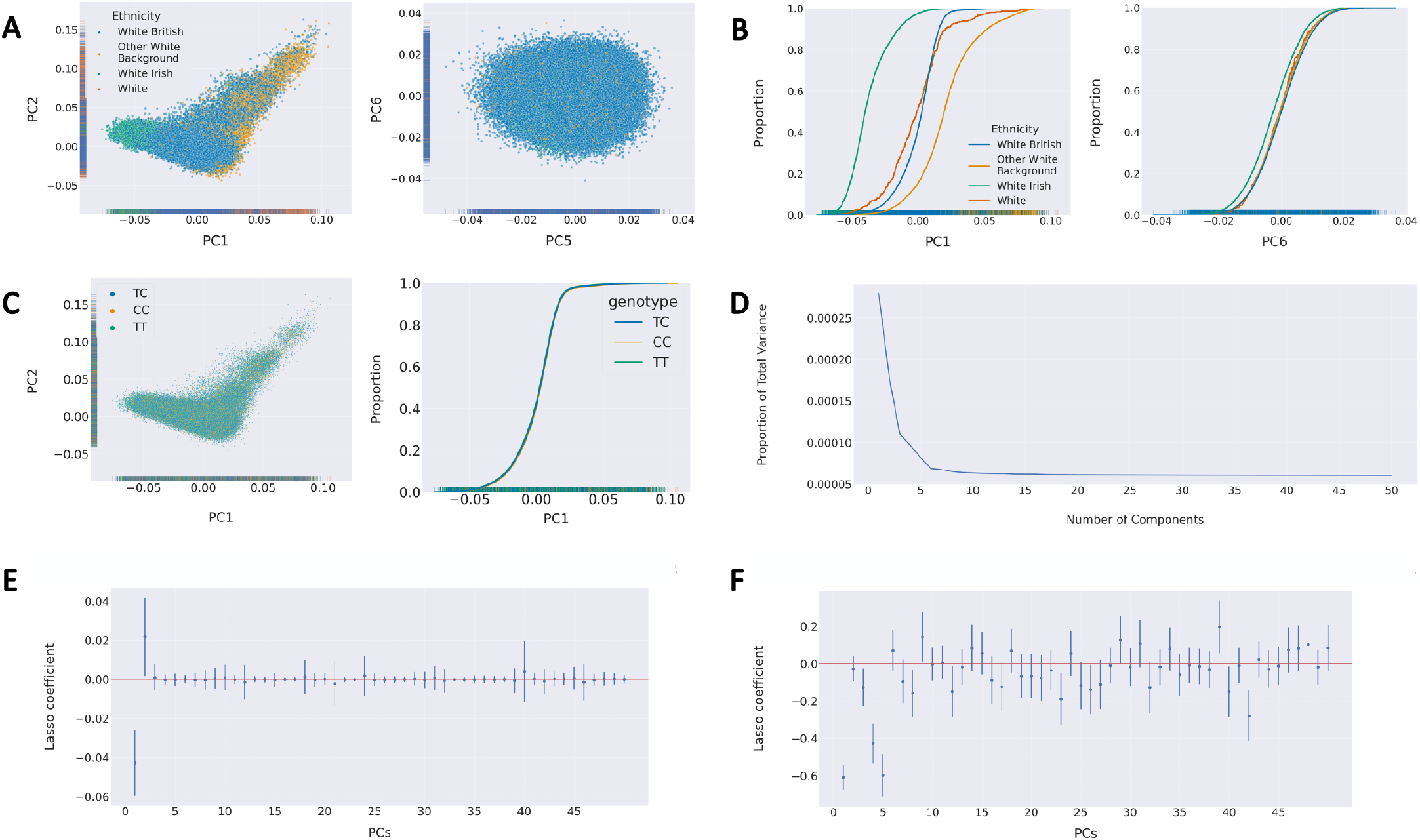
Population Structure within the UK Biobank cohort. (**A**) Principal component analysis labelled by ethnicity. Left: PC1 vs PC2 shows high level of population structure dependent on self-reported ethnicity. Right: PC13 vs PC14 shows a more symmetric shape suggesting that there is no ethnicity structure for PCs > 13. This is more clearly visible in (**B**) via the cumulative distribution analysis of ethnicity for PC1 and PC13. Left: In PC1 self-reported ethnicity populations have different distributions indicating that ethnicity drives the first PC. Right: In PC13 this separation has disappeared. For further cumulative distributions see Supplementary figures. (**C**) PC Analysis of genotypes within the cohort. Left: PC1 vs PC2 labelled with the three genotypes of the *FTO* intron variant, rs1421085. There is no evidence of population structure based on this genotype. Right: A cumulative distribution shows no separation between the distributions of the genotypes indicating that the variant is randomised in the cohort. When this is the case, population stratification (and hence the PCs) is not a confounder of the phenotype-genotype relationship and the only potential source of non-linearity in the functional dependence of phenotype on genotype and confounders is in the allelic copy (*Step 2A’* in Methods). (**D**) Scree plot shows that the proportion of variance explained by each additional PC plateaus after 6 to 8 PCs, when subset on ‘self-reported White’ UKB population, indicating that 8 PCs is sufficient to explain the population structure of this cohort. (**E**) and (**F**) Lasso regression results on 50 PCs, in which a statistically non-zero coefficient indicates that the corresponding PC component is relevant for the trait. These results show that the choice of PCs varies according to SNP-trait pair. The error bars represent two standard deviations from the mean derived from a bootstrap (*B* = 1000 bootstrap resamples) of the lasso coefficient. Any PCs whose error bars overlap with zero can be interpreted as adding no information to the population structure. (**E**) Lasso regression of trait “K76 other diseases of liver” (clinical_c_K76). This trait needs fewer than 5 PCs to explain the population structure within the cohort. (**F**) Lasso regression of trait “K20-K31 Diseases of esophagus, stomach and duodenum” (clinical_c_Block_K20-K31). This trait demonstrates dependence on PCs that fluctuates more extensively than the trait in panel E. In a bespoke analysis of a trait that shows fluctuation in the Lasso regression of trait on PCs in supernumerary components, we recommend including those higher PCs in the SL in addition to the result of the trait-independent analysis as part of a sensitivity analysis.

**Figure 6.**
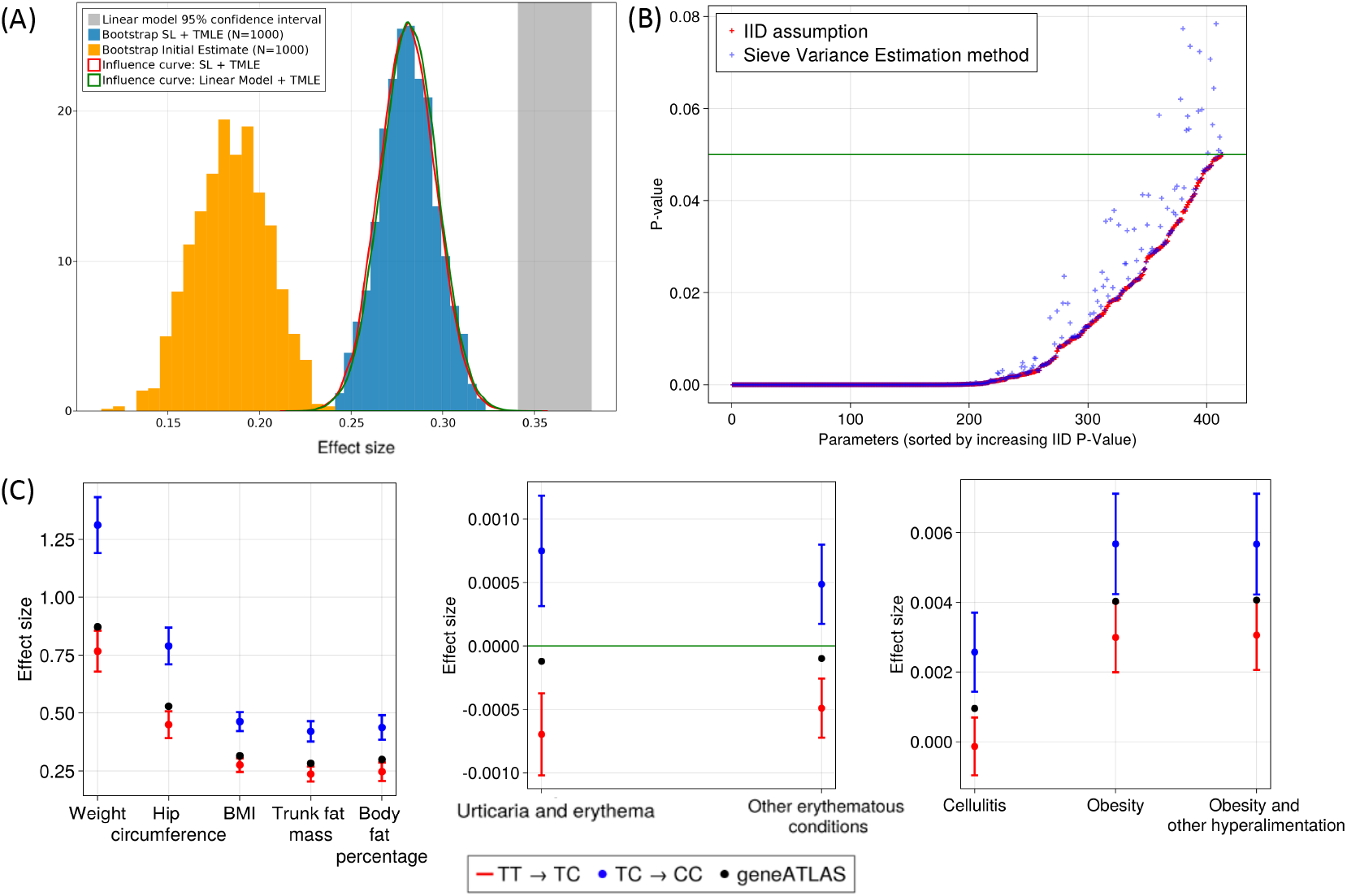
(**A**) **Inference results**. Comparison of various methods to estimate the effect size of rs1421085 on body mass index (BMI; X-axis; UK Biobank Data-Field 23104). The grey area indicates a 95% confidence interval on the effect size predicted by a linear-model. The orange histogram is the bootstrap distribution of the initial estimator when using a SL, without removing residual model-misspecification bias on the target quantity via TMLE. The blue histogram is also obtained from bootstrap but additionally using the TMLE. Finally, the red and green curves are the Gaussian distributions obtained via SL + TMLE and Linear Model + TMLE respectively. As can be seen, the effect sizes and corresponding p-values reported by the linear model alone, or SL alone, are overly optimistic or pessimistic, respectively. In both cases, the TMLE step brings the inference back to the same confidence region. See Panel A of Fig. 13 in Supplementary figures for a comparison between the initial estimates of effect sizes, relying on Super Learning, only against the final effect sizes after the TMLE step. Panel B of Fig. 13 in Supplementary figures shows a comparison between p-values reported by TarGene vs GeneATLAS. (**B**) **Sieve variance correction**. P-values obtained from two variance estimation methods for rs1421085. In red, the individuals in the UK Biobank are assumed to be i.i.d., while in blue, a sieve correction method is applied to account for the population dependence structure. Each p-value corresponds to a specific parameter of interest for which the initial i.i.d. estimate was under the 0.05 threshold. See Panel A of Fig. 14 in Supplementary figures for an example of a Sieve Plateau curve, and Panel B for the histogram of the genetic relationship matrix. (**C**) **Non-Linear effects**. A selection of traits for which rs1421085 TT → TC and TC → CC effect estimates are significantly different; Supplementary Table 2 contains the complete list. Two binary traits for which effect sizes have opposite sign (top), five continuous traits related to BMI (middle), and three further binary traits associated with BMI (bottom). Effect sizes are reported with associated 95% confidence intervals together with estimates from GeneATLAS’ LMM fits (black data points) (***Canela-Xandri et al., 2018***). The latter fall in-between our TT → TC and TC → CC estimates, indicative of an averaging effect.

##### Step 2A’

Non-confounding covariates need not be conditioned on for correct estimation In the estimation of the phenotype-genotype relation, covariates that do not confound this relation, *i.e*., that do not affect both phenotype and genotype, need not be taken into account. In the causal graph of Fig. 7, the left hand side represents a confounder *W* of the causal effect *V* → *Y* of genotype *V* on phenotype *Y*, whereas the right hand side *W* is not a confounder as it merely affects phenotype, not genotype. In other words, for the graph on the right hand side, the covariate *W* may be ignored when estimating the causal effect of *V* on *Y even though it affects outcome Y; however, the crux is that W does not affect the source V*. Mathematically, this means that, for any *υ*,

**Figure 7.**
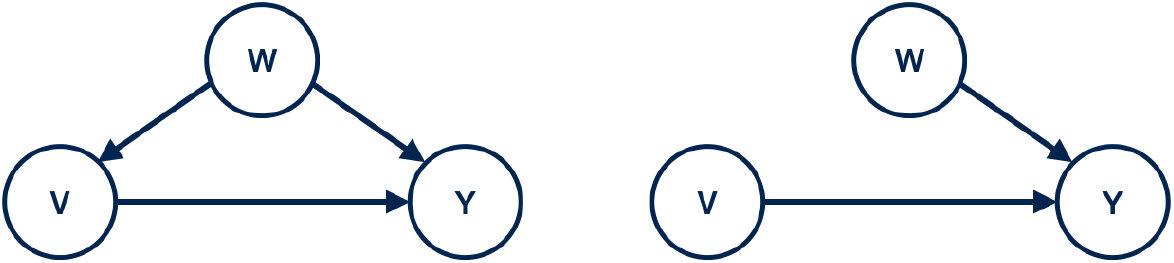
Two causal graphs from which we wish to identify the direct causal effect of *V* → *Y*. Left: *W* is a confounding variable which has to be correct for, *i.e*., conditioned on, for accurate causal identification. Right: *W* is not a confounder.

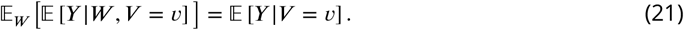

This can be deduced from the fact that the directed acyclic graph on the right hand side of Fig. 7 encodes the property *p*(*W* |*V*) = *p*(*W*) since *V* and *W* are independent. We have:

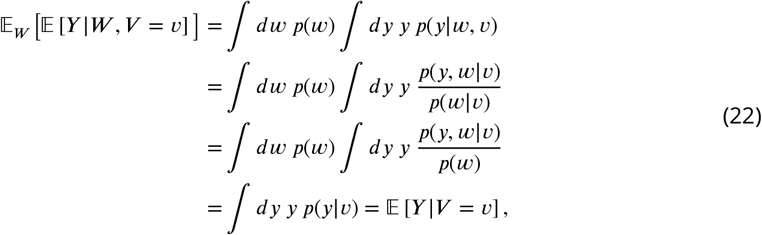

where in the second equality we have used the product rule of probabilities, in the third equality we have used the independence condition of *V* and *W*, and in the fourth equality we have exchanged 632 integrals and used ∫ *p*(*y*, *w*|*υ*)*dw* = *p*(*y*|*υ*).

#### Step 2B: TarGene leverages a diverse combination of algorithms via Super Learning

It is unnecessary to expend computational resources on estimating the full probability distribution *P* (*y*, *υ*, *w*) in order to evaluate the target parameter of Eq. 2. Indeed, only the part 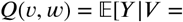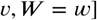 is required for estimating effect sizes per Eq. 7; similarly, the part 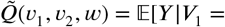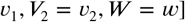 is required for estimating the interaction of Eq. 8. Super Learner (***van der Laan et al., 2007***) applies *k*-fold cross-validation to a diverse library of learning algorithms to obtain an estimate of the relevant part *Q*(*υ*, *w*) of *P*_0_, *i.e*., the expected phenotype given variant *υ* and confounders *w*; see Results and Fig. 3, Step 2. In *k*-fold cross validation, the data are split into *k* equally sized and disjoint folds. All algorithms are trained *k* times on the data, each time holding out a different ‘validation’ fold from the training procedure. The algorithms’ performance is subsequently validated on the held out fold. Finally, the best performing linear combination of algorithms is selected. The output of SL is an initial estimate 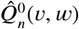 of the function *Q*(*υ*, *w*), as well as an initial estimate of the target parameter by plugging in:

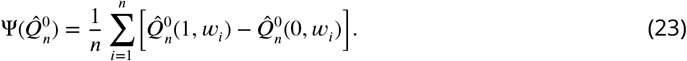

The average is taken over the population of size *n*, and *w*_*i*_ is the covariate of participant *i*.

In this paper, TarGene used the following SL specifications: (i) *k*-fold cross-validation or stratified *k*-fold cross-validation based on the outcome type (continuous or binary, respectively), here 3 ≤ *k* ≤ 20, selected adaptively based on the rarest class of each outcome (***Phillips et al., 2022***), and, (ii) included the constant fit, a regularized logistic/linear regression (ridge, *λ* = 1), a gradient-boosted tree (`n_round` = 20, default parameters otherwise), and HAL with hyper-parameters `max_degree` = 1, `smoothness_orders` = 1, `lambda` = 30 (***van der Laan, 2017***), as algorithms in the library; for full details see Practical Aspects. For confounding adjustment, we used the first 6 PCs from the genotypes, the age and the sex of each individuals. However, we note that for the optimal performance of HAL in more bespoke analyses, the parameter *λ*, tuning the total variation norm of the fit, should be left unspecified so that it is chosen by the algorithm’s internal cross-validation.

#### Step 3A: TarGene performs a targeted update via TMLE to remove bias

Although SL is optimised for estimating the function *Q*(*υ*, *w*), it is not optimised for estimating the DNA variant’s true effect size on phenotype, *i.e*., the target parameter Ψ(*P*_0_). As a result, there may be residual bias in the initial estimate Eq. 23, *i.e*., a discrepancy between the effect size estimate, 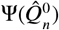, and its true value, Ψ(*P*_0_). Under mild assumptions, mathematical theory (see (***van der Laan, 2017***)) allows us to describe and analyse this discrepancy explicitly:

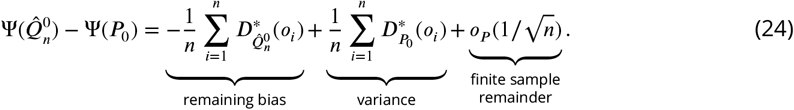

Here 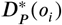 denotes the *efficient influence curve* of Ψ at *P* evaluated at the data point *o*_*i*_ = (*y*_*i*_, *υ*_*i*_, *w*_*i*_) of individual *i*. It quantifies the effect individual *i* has on the average effect size Ψ(*P*) across the population, *e.g*., a large effect if *i* is an outlier with a rare phenotype/genotype combination (the value of 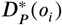 is large), and a small effect if *i* is a ‘typical’ participant (see Step 3B: Computing influence curves). The third term 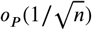 is due to the finite sample size of the data, and shrinks at rate 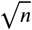 as sample size *n* increases.

Importantly, although the two averages on the right-hand side of Eq. 24 look similar, they play very different roles and have different interpretations. The first average is constructed from the initial estimate 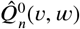 of *Q*(*υ*, *w*), and it quantifies the remaining bias due to the fit. We reduce this bias in order to optimise the bias-variance trade-off on the quantity of interest by applying Targeted Maximum Likelihood Estimation (TMLE, see (***van der Laan and Rose, 2011***)). Specifically, applying TMLE updates the initial fit 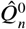 repeatedly until, at the final iteration denoted 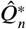, the remaining bias term in Eq. 24 is approximately zero, namely 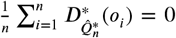; in practice, a remaining bias term of 10^−16^ to 10^−7^ is numerically observed depending on t*n*he target quantity of interest. For the effect size and interaction target quantities discussed in this work, it is a mathematical fact that convergence takes place in a single step so that no iteration is necessary. The final TL estimate of the DNA variant’s effect size on phenotype is

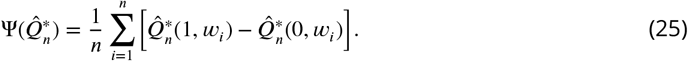

In contrast, the second average in Eq. 24, containing terms 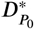, depends on the ground truth probability distribution, *P*_0_, and cannot be changed by an improved analysis (apart from increasing the data size). It is responsible for the variance on the estimate 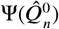. Indeed, after removing the remaining bias term via TMLE, Eq. 24 implies

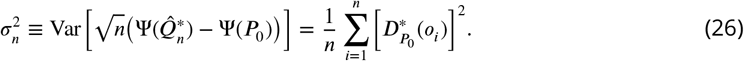

The right-hand side of this equation can be directly estimated from data and quantifies the variance on the final effect size estimate 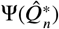. Furthermore, it allows for the direct construction of an approximate Wald-type 95% confidence interval:

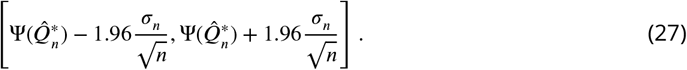

This approach is valid since the TMLE step and Eq. 24 (with vanishing first term on the right-hand side) imply that 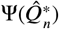 is normally distributed as *n* becomes large, provided the third term in Eq. 24 is indeed of order 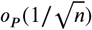. The latter condition holds as long as the product of *Q*_0_ and *g*_0_ is estimated at a rate of *n*^−1/2^ because the effect size and interaction target quantities have the *double robust property*; see Step 3D: TL estimates are double-robust for details. Finally, we can directly construct a p-value on the estimate of effect size from this confidence interval.

#### Step 3B: Computing influence curves

An asymptotically linear estimator behaves, for large sample sizes, like an average of independent identically distributed random variables. These random variables, called *influence curves*, are functions of the data. The Central Limit Theorem can be used to analyse the variance of asymptotically linear estimators. More precisely, an estimator 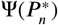 of a quantity of interest is asymptotically linear if

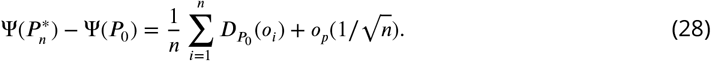

Here 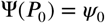 is the true value of the parameter, 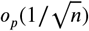 is a finite-sample term that shrinks to zero at rate 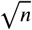 as sample size *n* increases, and 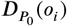 is the influence curve of Ψ at the probability distribution *P*_0_ evaluated at the *i*th data point *o*_*i*_. The expectation of 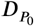 with respect to the ground truth *P*_0_ is zero, *i.e*., 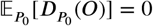.

The influence curve of the average treatment effect Ψ_1_(*P*) of Eq. 7 can be found in (***van der Laan and Rose, 2011***, App. A.3). It is the following function of data *O* = (*Y*, *V*, *W*):

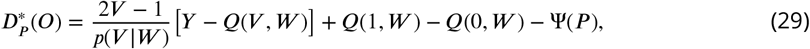

where 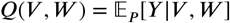. Here *H*(*V*, *W*) = (2*V* − 1)/*p*(*V*|*W*) is the ‘clever covariate’.

Since the target parameter for interaction 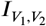 of Eq. 8 has recently been introduced in (***Beentjes and Khamseh, 2020***), we derive its influence curve here for the first time. We record our result as

##### Proposition 0.1

*Let* 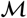 *be a non-parametric statistical model containing probability distributions of O* = (*Y*, *V*_1_, *V*_2_, *W*) *where Y is any outcome*, *V*_1_ *and V*_2_ *are binary or categorical variables*, *and W is any covariate*. *Let* (*a*, *b*) *be categories of* (*V*_1_, *V*_2_), *and consider the target parameter*

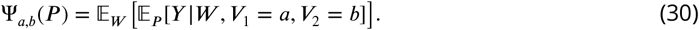

*The efficient influence curve of* Ψ_*a*,*b*_ *is given by*

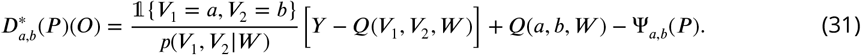

*Here we have defined the function* 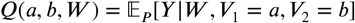.

Since the influence curve of a sum of target parameters is equal to the sum of their influence curves, we immediately deduce the influence of interaction 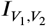:

##### Corollary 0.2

*The influence curve of* 2-*point interaction* 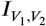 *of Eq. 8 equals*

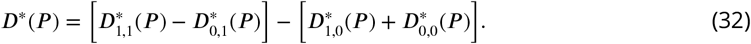

We prove Proposition 0.1 in Appendix A: Proofs.

#### Step 3C: TMLE updates the initial fit to obtain mathematical guarantees

The targeted update step in TMLE removes the remaining bias term in Eq. 24 in order to optimise the bias-variance trade-off on the quantity of interest, thus making the effect size estimator asymptotically normal and allowing for the construction of Wald-type 95% confidence intervals. This step proceeds by fluctuating the initial SL fit 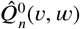 of the conditional phenotype 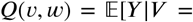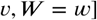 in the direction of the efficient influence curve 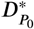 of the quantity of interest, *e.g*., the effect size of Eq. 29 or genotype-genotype interaction of Eq. 31. This fluctuation consists of a simple one-dimensional maximum likelihood estimation (MLE) of a real-valued parameter *ϵ* in an auxiliary statistical model. Specifically, for binary phenotypes, the fluctuation is a logistic regression,

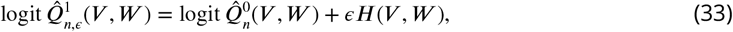

so that the property 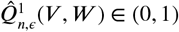 is preserved. For continuous phenotypes, the fluctuation is a linear regression (hence with a normally distributed noise term),

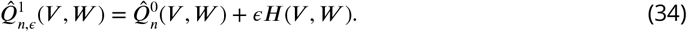

In both cases, the initial SL fit 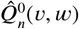 is taken as off-set and the coefficient *ϵ* in front of *H*(*V*, *W*) is estimated; the fitted value of *ϵ* in the first update is denoted by 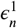. The clever covariate depends on the quantity of interest, satisfying *H*(*V*, *W*) = (2*V* − 1)/*p*(*V*|*W*) for effect sizes and *H*(*V*_1_, *V*_2_, *W*) = (2*V*_1_ − 1)(2*V*_2_ − 1)/*p*(*V*_1_, *V*_2_|*W*) for variant-variant interactions. Performing weighted, instead of standard, logistic (or linear) regression with weight the reciprocal of the propensity score and clever covariate *H*′(*V*, *W*) = 2*V* − 1 for effect sizes and *H*′(*υ*_1_, *V*_2_, *W*) = (2*V*_1_ − 1)(2*V*_2_ − 1) for interactions, results in more robust estimates when near positivity violations are present. We fit the treatment mechanism *g*(*υ*, *w*) = *p*(*υ* = *v*|*W* = *w*) (for effect sizes) and *g*(*υ*_1_, *υ*_2_, *w*) = *p*(*V*_1_ = *υ*_1_, *V*_2_ = *υ*_2_|*W* = *w*) (for interactions) using SL (with the same specification as reported for 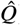), and denote the corresponding fit by 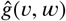. The covariates *W* used are the principal components, age and sex of the participants. Performing MLE means solving the score equation,

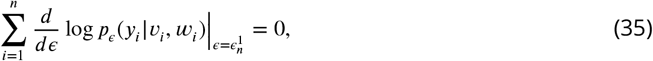

where *p*_*ϵ*_(*y*|*υ*, *w*) is the probability density of phenotype *y* given genotype and covariates *υ*, *w*. By construction of both fluctuations, Eq. 35 evaluated at *ϵ* = 0 equals the empirical mean of the efficient influence curve. Thus, we iterate the TMLE step, each time taking the updated fit 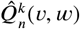 as off-set, until the fitted parameter 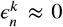. Since the clever covariate is independent of *Q*, it is a mathematical fact that this algorithm converges in a single step, that is, 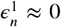 and 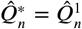. Then Eq. 35 reads

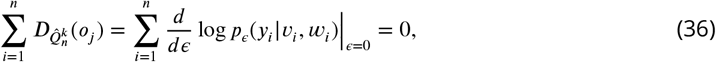

and we have updated 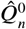 to 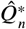 ∶= 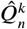 so as to eliminate the remaining bias term in Eq. 24, thus making the estimate 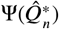 asymptotically normal with optimised bias-variance trade-off.

#### Step 3D: TL estimates are double-robust

In order to obtain the final estimate, 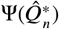, of a target parameter such as effect size or variant-variant interactions, the TL workflow requires the estimation of two quantities: the true conditional phenotype, *Q*_0_, and the true treatment mechanism, *g*_0_. This is an advantageous feature since TL estimates are double-robust in the sense that 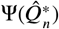 is a consistent estimate of the ground truth Ψ(*P*_0_) provided either (or both) *Q*_0_ and *g*_0_ are estimated consistently at a rate of convergence faster than *n*^−1/4^. Put differently, if only one of *Q*_0_ and *g*_0_ is estimated *in*correctly, *e.g*., by a misspecified model, then the TL estimate is nevertheless consistent. If both *Q*_0_ and *g*_0_ are estimated consistently at a product of rates faster of at least *n*^−1/2^, then 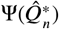 is an efficient estimator of Ψ(*P*_0_), *i.e*., the estimator with the smallest variance in its class.

The double-robustness property of a TL estimator is established separately for each target parameter. Here, we first recall the double-robustness of effect size to then establish that the interaction target parameter of Eq. 8, introduced in (***Beentjes and Khamseh, 2020***), is double-robust as well. The general approach is to recall the first-order approximation of the target parameter at a probability distribution *P* in terms of the influence curve as in Eq. 28,

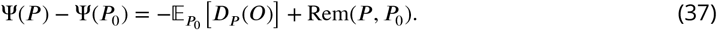

in terms of its influence curve, and analyse the second-order remainder term,

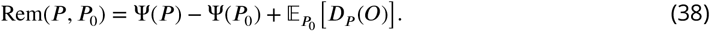

For the parameter Ψ_1_ of Eq. 2, measuring the effect size of a DNA variant *V* on a phenotype *Y* correcting for confounders *W*, we have 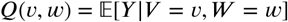 and *g*(*υ*, *w*) = *p*(*V* = *υ*|*W* = *w*). It is well known that the remainder term of Ψ_1_ satisfies

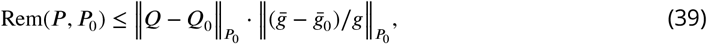

where 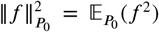 for a function *f* of the data *O* = (*Y*, *V*, *W*), we write 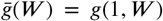 and 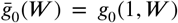. This inequality is a special case of the Cauchy–Schwartz inequality. We show in Appendix A: Proofs that the same inequality holds for the second-order exact remainder of the interaction target parameter of Eq. 8.

##### In practice

The above description of the estimation framework of Targeted Learning is predicated on the estimation of the effect size of a single SNP on a single phenotype measured on independent and identically distributed data. However, if the data are *dependent*, care must be taken in estimating the variance on the estimates; in particular, Eq. 26 needs to be generalised, see Eq. 49. Furthermore, if multiple effect sizes are estimated, multiple hypothesis correction must be incorporated in order to bound type I errors, such as FDR control, and obtain joint p-values. Both of these further steps are required when dealing with large-scale population genetics data, such as the UKB.

### TarGene identifies non-linear effects of allelic copies on phenotype

TarGene estimates both effect sizes Ψ_1_(*P*) and Ψ_2_(*P*) in Eq. 2 and Eq. 6 separately and can, thus, be leveraged to (i) determine significant non-linear effects of allelic copies on phenotypes as well as (ii) classify the type of SNPs and/or phenotypes for which such non-linearities occur. In practice, TarGene does this by combining the asymptotic description of both TL estimates as averages of independent random variables with the Central Limit Theorem (CLT). More precisely, let 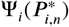 be the *i*th effect size with efficient influence curve 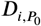 for *i* = 1, 2. By TL theory, 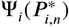 is the empirical average over influence curves and thus, by the CLT, asymptotically normally distributed:

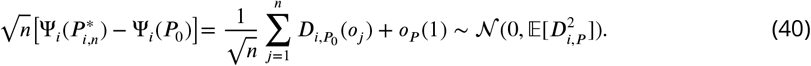

Here, we are interested in the difference target parameter Ψ_Δ_(*P*) = Ψ_2_(*P*) −Ψ_1_(*P*). Taking the difference of the quantities in Eq. 40 for *i* = 1, 2 yields a description of the difference target parameter minus the difference ground truth as an average over influence curves:

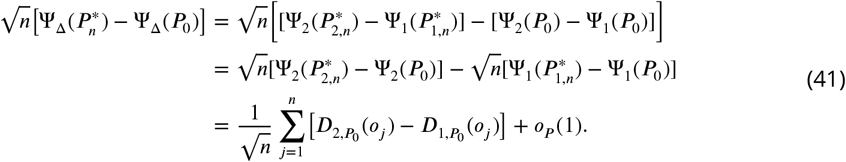

The asymptotic normal distribution of 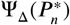 and its variance again follow from the CLT, the application of which is a special case of the functional delta method:

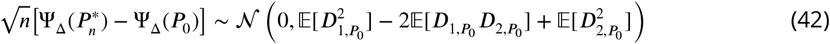

In practice, the variance on the right-hand side is estimated by replacing the influence curves by their sample averages, followed by an update to accurately account for the population dependence structure in the cohort (see Step 4 below and Eq. 50).

#### Step 4: TarGene accounts for population dependence structure

Biobank cohorts consist of participants who are, to some extent, related due to ancestry or kinship. TarGene accounts for this population dependence structure by appropriately adjusting variance estimates of effect sizes and interactions via Sieve Plateau (SP) variance estimators (***Davies and van der Laan, 2014***) which, in turn, are based on the genetic distance between participants as encoded in the genetic relationship matrix.

##### Genetic Relationship Matrix

Many statistical analyses rely on the assumption that the data are *independently* sampled from the population; it is also often required that the data be *identically distributed* but this assumption is typically not essential. However, this independence assumption no longer holds for the participants in the UK Biobank since many of whom are, to some extent, genetically related. Such genetic similarity can occur on a sub-population level due to ancestry (*e.g*., being white Irish), or on an individual level due to kinship (*e.g*., parents, children, cousins). Moreover, genetically similar individuals may share diet and environment inducing further dependence (***Abdellaoui et al., 2022***).

The genetic similarity of two individuals *i* and *j* is quantified by the sample correlation coefficient *G*_*ij*_ between their (centred and scaled) SNPs. Together, these coefficients form the *Genetic Relationship Matrix* (GRM), denoted *G*, of size *N* × *N* where *N* is the number of individuals in the population. More precisely, given a set of *R* SNPs, we have

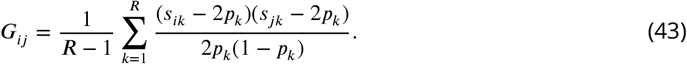

Here *s*_*ik*_ ∈ {0, 1, 2} denotes the number of copies of the reference allele for individual *i* at SNP *k*, and *p*_*k*_ ∈ (0, 1) denotes the frequency of the reference allele at SNP *k* over the population of *N* individuals. In particular, the population average of *s*_*ik*_ equals twice the reference allele frequency at SNP k, *i.e*., 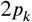 (one for each strand copy), so

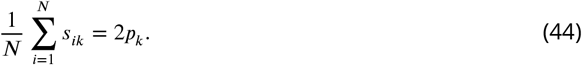

Thus 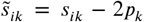 is the zero-centred count of the number of copies of the reference allele of individual *i* at SNP k. Considered as a random variable, 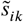 takes on three values. Assuming reference alleles are sampled binomially with mean frequency *p*_*k*_, the standard deviation of 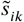 equals 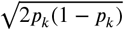. This explains the additional factor in Eq. 43 that scales the variables 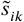 and 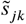 so as to have unit variance. Finally, note that the GRM depends on the set of *R* selected SNPs. These SNPs should be chosen amongst genotyped (not imputed) SNPs that, in addition, are not in LD with one another.

##### Sieve Plateau Variance Estimators

In TarGene, we neither assume individuals are independent nor do we impose the strong restrictive assumptions of an LMM. Instead, we incorporate the genetic dependence of individuals model-independently in our Targeted Learning framework described in the section TarGene provides mathematical guarantees and realistic p-values. This approach, drawn from (***Davies and van der Laan, 2014***), is based on mathematical theory and addresses all the above issues. It generalises Eq. 26 for the variance of the effect size target parameter, 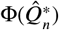, by constructing *Sieve Plateau (SP) variance estimators* that incorporate genetic dependence of individuals. These estimators result in valid confidence intervals and, ultimately, realistic and valid p-values having correctly accounted for population stratification.

We now illustrate how data dependence impacts the variance estimate of Eq. 26. Since individuals *i* and *j* are in general dependent, their data *O*_*i*_ = (*Y*_*i*_, *V*_*i*_, *W*_*i*_) and *O*_*j*_ are also in general dependent. As a result, the same holds for their corresponding influence curves 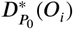 and 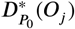. The problem arises since for two random variables *X*_1_ and *X*_2_ the variance of their sum is *not* in general equal to the sum of their variances. The difference is exactly twice the covariance of *X*_1_ and *X*_2_,

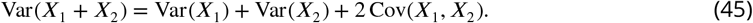

The impact of this difference may be large depending on the size of the covariance Cov(*X*_1_, *X*_2_); note that the covariance of two random variables vanishes when they are independent. Thus, rather than Eq. 26, the true variance on 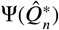 is given by

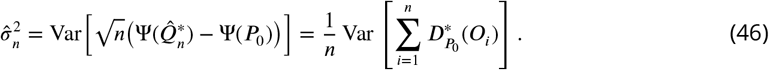

Now, the distinction between Eq. 26 and Eq. 46 is only relevant for significantly genetically similar individuals. SP variance estimators define a cut-off *τ* for the allowed genetic distance between individuals, and set the covariance to zero between individuals that are sufficiently genetically dissimilar. We obtain a variance estimate, 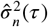, for each value of *τ*. The true variance of the estimate is obtained where the function 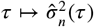 plateaus.

To construct SP variance estimators, we proceed as follows:

1. Using the GRM, we define a *genetic distance* between individuals *i* and *j* as

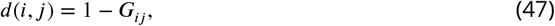

where *G*_*ij*_ is the sample correlation coefficient of Eq. 43 quantifying the genetic dependence between individuals *i* and *j*. Since correlation is bounded, |*G*_*ij*_| ≤ 1, the genetic distance is non-negative and never larger than two, *i.e*., 0 ≤ *d*(*i*, *j*) ≤ 2. Biologically, if two individuals *i* and *j* have identical SNPs they are fully correlated, *G*_*ij*_ = +1, and thus have zero genetic distance, *d*(*i*, *j*) = 0, as expected.
2. Given a value for the cut-off *τ* ∈ [0, 1], we define a SP variance estimator as

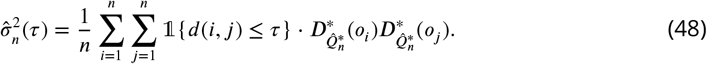 Here, the term 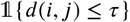 equals 1 if the genetic distance between individuals *i* and *j* is at most *τ*, *i.e*., *d*(*i*, *j*) ≤ *τ*, and it equals 0 otherwise. The biological interpretation of these estimators is as follows. The correlation between the influence curves of individuals *i* and *j*, estimated by the term 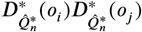, is taken into account only if the genetic distance between individuals *i* and *j* is at most *τ*. Thus, the SP variance estimator 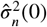, *i.e*., when *τ* = 0, assumes all individuals are *independent*. By increasing *τ*, we first take the covariance between strongly genetically dependent individuals into account for low *τ*, and then incorporate the covariance of more weakly dependent individuals as *τ* increases up to *τ* = 1.
3. We construct the variance estimator 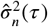 for a number of values of the cut-off *τ*, *e.g*., 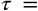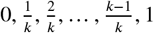. Then we fit the function 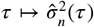 and select the value of *τ* where the function plateaus, call it *τ*_0_.
4. The correct variance estimate is then 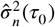.

Under mild assumptions (***Davies and van der Laan, 2014***, Theorem 1), the distribution of the effect size estimate 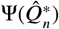 of SNP *V* on phenotype *Y* (or the effect of an epistatic interaction on phenotype) is asymptotically normal, and the SP variance estimator allows for the construction of an approximate 95% Wald-type confidence interval for the estimate 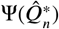 in the usual way, namely

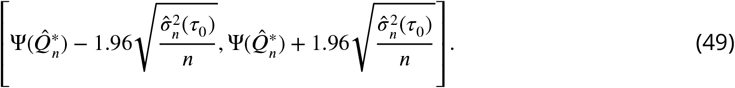

From here, we obtain realistic p-values correctly accounting for population dependence.

Similarly, we need to take into account population dependence in order to obtain a realistic estimate of the variance on 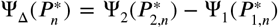 in Eq. 42. If this difference of effect sizes of a DNA variant *V* on a trait *Y* is significant, the effect of an additional allelic copy is non-linear. We construct an SP estimator for the variance on this difference by following steps (1)–(4) above, with the exception of appropriately generalising Eq. 48 in step (2) as follows. Given a value for the cut-off *τ* ∈ [0, 1], the estimator is

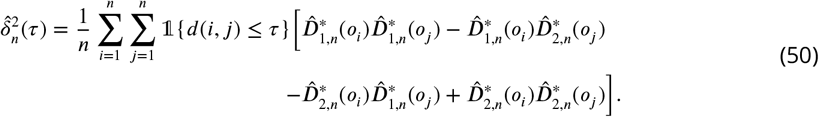

Here we have used the short-hand 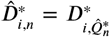 for the influence curve of effect size *i* evaluated at the final TMLE estimate 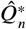 of the conditional expectation 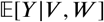.

#### Step 5: TarGene controls for multiple hypothesis testing

In answering a typical question in population genetics, multiple hypotheses are tested simultaneously. Depending on the nature of the question, a specific multiple hypothesis correction is necessary to bound the error rate of interest. Error rates commonly employed are (i) the probability of at least one false discovery, or *family-wise error rate*, FWER = *P* (*V*_*n*_ > 0), and (ii) the expected ratio of false discoveries to true discoveries, or *false discovery rate*, 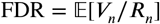. Here *n* denotes the sample size, *R*_*n*_ denotes the (known) number of discoveries, and *V*_*n*_ denotes the (unknown) false discoveries or Type I errors; see Fig. 3E. Once an error rate and bound has been chosen, *e.g*., one seeks to bound the FDR at ≤ 0.05, then a multiple hypothesis correction procedure is to be chosen. The better procedure is the one that minimises false negatives (Type II errors, denoted by *T*_*n*_ in Fig. 3E), *i.e*., maximises power, at the given Type I error control.

The literature on multiple testing procedures is extensive, see for example (***Dudoit and van der Laan, 2008***). Procedures differ mainly in that (i) they depend on the marginal distribution of the test statistics only (*marginal* procedure) or on their joint distribution (*joint* procedure), (ii) the rejection criteria of the next test is independent of the outcome of the previous tests (*single-step* procedure, such as Bonferroni correction) or the rejection criteria depend on the outcome of previous tests (*step-down* procedure, *e.g*., Benjamini–Hochberg). Further considerations can be taken into account, all of which are aimed at maximising power at a given error rate whilst respecting the dependence structure of the tests’ null hypotheses. For example, a computationally more intensive joint procedure is unwarranted when test statistics are largely independent.

Since TarGene produces asymptotically normal estimators which are empirical means of their efficient influence curve, a vector of these estimates similarly equals an empirical mean of the vector of efficient influence curves. By the multi-variate Central Limit Theorem, their joint distribution is then again multi-variate normal. Thus, when simultaneously testing for significance of multiple (i) effect sizes, (ii) non-linearity of effect size, and/or (iii) epistatic or gene-environment interactions, the asymptotic joint distribution of the corresponding null distribution is known. As a consequence, researchers can take advantage of both marginal and joint procedures to maximise power whilst bounding their desired Type I error rate.

In this work, we use the marginal step-down Benjamini–Hochberg procedure of (***Benjamini and Hochberg, 1995***) to control the FDR at ≤ 0.05.

##### Practical Aspects

To enable reproducible research and future usage, we provide [TarGene](https://github.com/TARGENE/targene-pipeline) as a Nextflow pipeline. For more specific analyses, even outside the genetic field, we also release a general purpose [Julia](https://julialang.org/) package for Targeted Minimum Loss-Based Estimation at TMLE.jl.

Our proposed method is computationally intensive by nature. However, performance is critical for routinely conducted large scale genetic studies like PheWAS or GWAS. We show below that, access to modern computing resources, makes TarGene possible in those large scale settings. In our case, all runs were performed on the Edinburgh high-performance [Eddie](https://www.ed.ac.uk/information-services/research-support/research-computing/ecdf/high-performance-computing) cluster. The most computationally intensive processes in TarGene are (i) the TMLE process, including nuisance parameters estimation (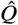 and 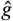) and the targeting step, and (ii) the Sieve Variance Plateau estimation process. Below, we present run-times for both processes.

###### TMLE Process

In this section we investigate the run time of the TMLE process for the two most common genetic studies: GWAS and PheWAS. In both cases we are thus computing the Average Treatment Effect for each individual variant on trait by comparing the major/minor to the major/major genotype. Covariates were set to include the first 6 principal components, age and sex. Finally, the benchmark is performed on a 8-core compute node representative of a modern laptop. The code to reproduce the experiment is available here.

We investigate the 4 following nuisance parameters estimation strategies (for both 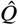 and 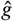) from the most naive to the most comprehensive:

- GLM: Standard generalized linear model
- GLMNet: GLM with regularization hyperparameter tuning over 3-folds cross-validation.
- XGBoost: The famous [gradient boosting trees](https://xgboost.readthedocs.io/en/stable/) method with hyperparameter tuning over 10 different settings in a 3-folds cross-validation scheme.
- SL: Super Learning including both the previous XGBoost and GLMNet combined with an outer 3-folds cross-validation.

We first focus on the PheWAS setting for which run time estimates are provided in Table 1. It is important to emphasise that in a typical PheWAS the estimation of the propensity score, 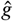, needs only to be performed once and can be re-used for all traits across the study. The computational complexity is thus driven by the estimation of each regression, 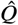, and associated targeted steps. We also note that the same remark holds for the targeting steps corresponding to the various genetic changes. Computing the effects of the additional major/minor → minor/minor and major/major → minor/minor would only cost two additional targeting steps while re-using the current 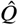. In all cases, running a PheWAS using TarGene is possible even without access to a high-performance computing platform.

**Table 1.** PheWAS run times for various nuisance parameters estimation strategies.

| Learning Algorithm | Time (hours) |
| --- | --- |
| GLM | 2.2 |
| GLMNet | 4.5 |
| XGBoost | 8.8 |
| SL | 30 |

We now turn to the GWAS setting for which run time estimates are provided in Table 2. In this case, 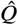 and 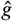 need to be estimated for each variant. In order to obtain a run time estimate for a GWAS it is thus sufficient to compute the run time for one variant and simply multiply by the number of variant in a study (We assume 600 000). Because we noticed the run time of the propensity score fit varies depending on the variant, we instead run the TMLE process over 100 variants and report the mean runtime as a more accurate estimate. While it would be impossible to run a GWAS on a personal laptop, we find that access to a modern computing platform makes this kind of study feasible using TarGene.

**Table 2.**
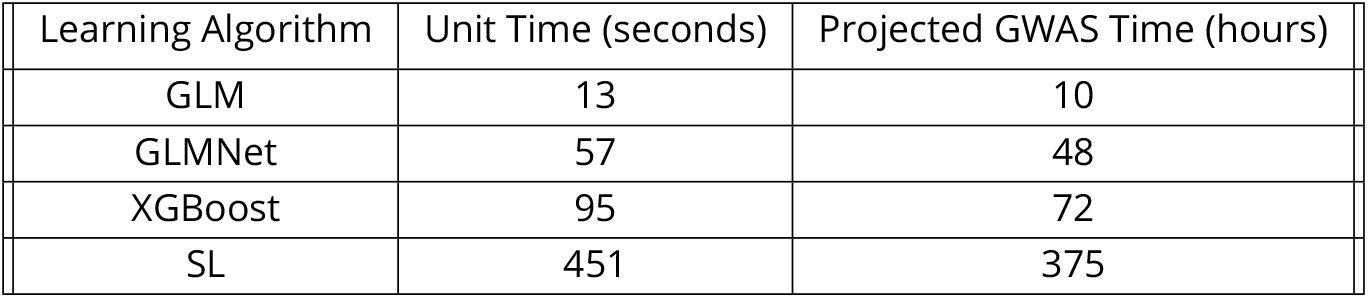
GWAS run times. The unit time corresponds to a single variant/trait pair. The projected GWAS time assumes 600 000 variants and 200 folds parallelization.

| Learning Algorithm | Unit Time (seconds) | Projected GWAS Time (hours) |
| --- | --- | --- |
| GLM | 13 | 10 |
| GLMNet | 57 | 48 |
| XGBoost | 95 | 72 |
| SL | 451 | 375 |

###### Sieve Variance Plateau Process

We next turn to the Sieve Variance Plateau estimation for which benchmarks on a 20-core compute node are presented in Table 3. For each parameter, the procedure computes a given number of variance estimates, here either 10 or 100 *τ*-values. As can be seen from the table, computational time increases sub-linearly with both the number of estimates and the number of parameters. This is mainly because, for a small number of parameters, the computational time is driven by reading the GRM from disk hence under-using the multi-threading power of the node. As soon as the number of parameters becomes large, *e.g*., for a GWAS or PheWAS, all cores can be utilised simultaneously to maximum efficiency. We thus recommend grouping together those operations as much as possible.

**Table 3.** Sieve Plateau variance estimation benchmarks.

| Number of estimates per curve | Number of parameters | Time (s) | Time (h) |
| --- | --- | --- | --- |
| 100 | 419 | 71836 | 19.9 |
| 10 | 419 | 11926 | 3.3 |
| 100 | 1 | 30966 | 8.6 |
| 10 | 1 | 6779 | 1.8 |

Finally, we note that in our experience on UKB, a nation-wide cohort with relatively low levels of relatedness, genetic dependence is less likely to affect the estimates and their variance, and so the SP variance estimator is unlikely to alter the significance of results. We therefore recommend, in case computational resources are limited, that the SP variance estimator be run for estimates just below the required significance threshold. In contrast, for cohorts displaying strong relatedness, *e.g*., island cohorts, a model-free dependence correction such as the SP variance estimator may have a more prominent role. Typically, such cohorts are orders of magnitude smaller in size than UKB, which greatly reduces memory requirements and computational run time.

## Acknowledgements

This research has been conducted using the UK Biobank Resource under Application Number 53116. MvdL is supported by NIH grant R01AI074345. CPP is funded by the MRC (`MC_UU_00007/15`). AK was supported by the XDF Programme from the University of Edinburgh and Medical Research Council (`MC_UU_00009/2`).

## Appendix A Proofs

In this section, we prove Proposition 0.1. Below, we recall the content of this statement for the reader’s convenience. As a direct corollary, this results in the computation of the influence curve of the interaction target parameter of Eq. 8, required for TMLE.

## Proposition 0.3

*Let* 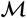 *be a non-parametric statistical model containing probability distributions of O* = (*Y*, *V*_1_, *V*_2_, *W*) *where Y is any outcome*, *V*_1_ *and V*_2_ *are binary or categorical variables*, *and W is any covariate*. *Let* (*a*, *b*) *be categories of* (*V*_1_, *V*_2_), *and consider the target parameter*

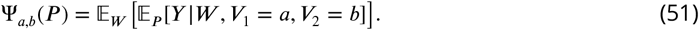

*The efficient influence curve of* Ψ_*a*,*b*_ *is given by*

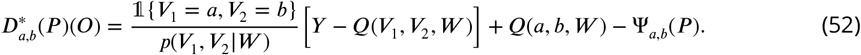

*Here we have defined the function* 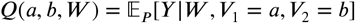.

Since the influence curve of a sum of target parameters is equal to the sum of their influence curves, we immediately deduce the influence of interaction 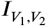:

## Corollary 0.4

*The influence curve of* 2-*point interaction* 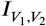 *of Eq. 8 equals*

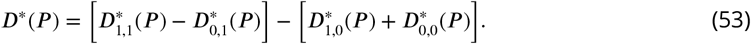

A proof of a similar result, from which this Proposition can be derived via the delta method, can be found in, *e.g*., Appendix A3 of (***van der Laan and Rose, 2011***). For the sake of completeness, we include a proof here.

*Proof*. The probability density function of *P* factors as

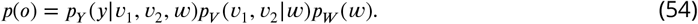

Let *P*_*ϵ*_ be any path in 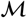 through *P* at *ϵ* = 0 with score *S*(*o*) = (*d*/*dϵ*)|_*ϵ*=0_ log *p*_*ϵ*_ (*o*), where *p*_*ϵ*_ is the probability density function of *P*_*ϵ*_ with respect to *λ*. By Theorem 3.2 of (***Tsiatis, 2006***), a gradient *D*_*a*,*b*_(*P*) of Ψ_*a*,*b*_(*P*) can be obtained by expressing the path-wise derivative of Ψ_*a*,*b*_(*P*_*ϵ*_) as

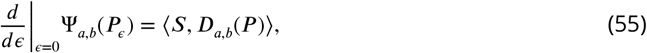

where the inner product ⟨−, −⟩ on the right-hand side is taken in the Hilbert space 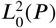 of mean-zero functions that are square-integrable with respect to *P*.

We now compute the path-wise derivative of the target parameter. First, we find

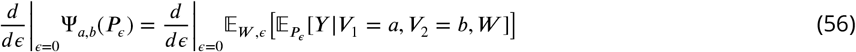

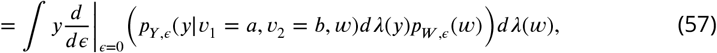

where derivative and integral can be exchanged by Lebesgue’s dominated convergence theorem. Next, since *V*_1_, *V*_2_ can be considered as binary random variables, we have

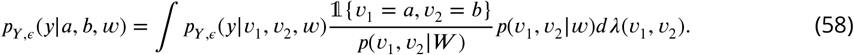

Here 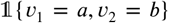 equals 1 when both *υ*_1_ = *a* and *υ*_2_ = *b* and vanishes otherwise. By another application of the dominated convergence theorem, we obtain

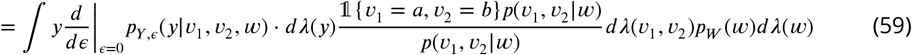

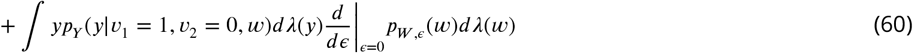

We consider these two integrals separately, utilising the Hoeffding decomposition; see, *e.g*., §11.4 of (***van der Vaart, 1998***). For the second integral, by the ordering *O* = (*Y*, *V*_1_, *V*_2_, *W*), this results in

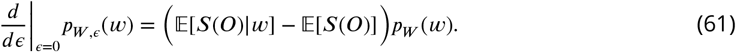

We now compute the second integral *I*_2_ to be

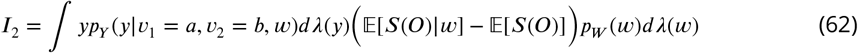

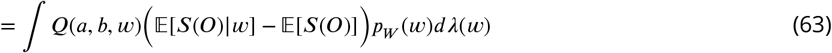

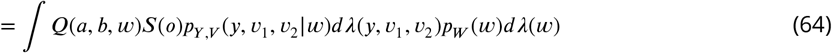

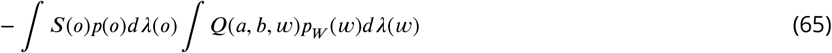

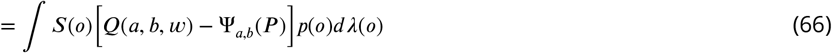

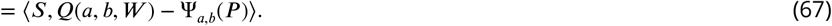

For the first integral, again applying the Hoeffding decomposition yields

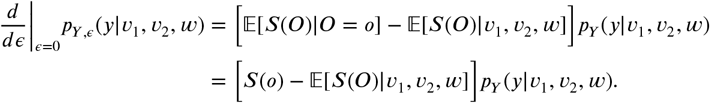

Next, we compute the first integral *I*_1_, which immediately splits into two pieces:

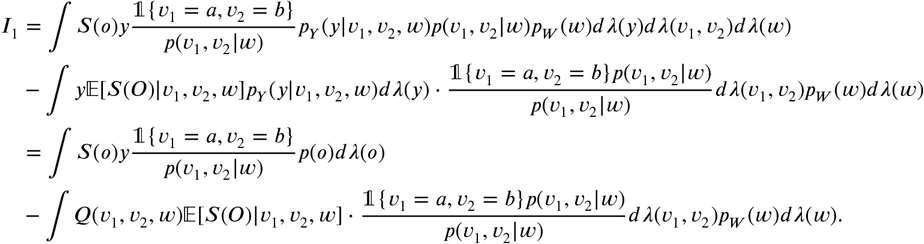

The first piece is of the required form. In the second piece 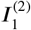 of integral *I*_1_ we have integrated out the *y*-dependence, which results in the factor 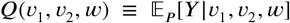. Next, we write 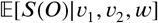 as an integral to have *p*(*o*) under the integral:

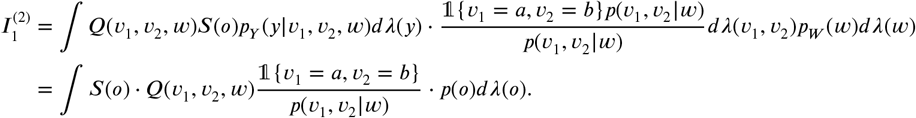

We infer that

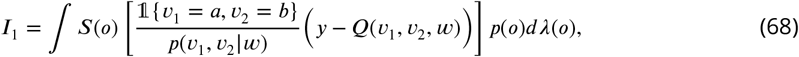

and conclude that the influence curve of Ψ_*a*,*b*_(*P*) is given by

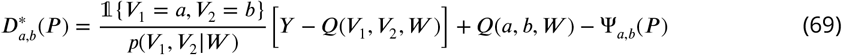

as claimed.

Given the influence curve of 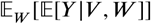, we can show the double-robustness of any target parameter that is a linear combination of such terms. This holds because both the exact remainder,

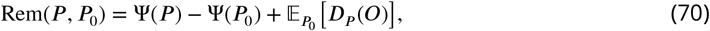

and the influence curve of a linear combination of target parameters equals the linear combination of exact remainders and influence curves respectively.

## Proposition 0.5

*Let* 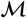 *be a non-parametric statistical model containing probability distributions of O* = (*Y*, *V*, *W*) *where *Y* is any outcome (e.g., trait), *V* a binary or categorical variable (e.g., a DNA variant), and *W* is any covariate. Consider the target parameter*

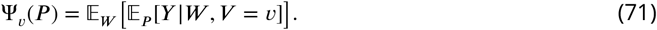

*The exact remainder of* Ψ_*υ*_ *is given by*

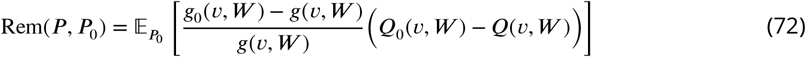

We explain in what sense the TMLE of ATE or interaction is double robust. Given a function *f* of the data *O* = (*Y*, *T*, *W*), we write 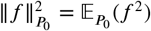 for its *L*^2^-norm. This norm is induced by the inner product 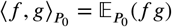 where *f*, *g* are two square-integrable functions of the data *O* = (*Y*, *T*, *W*). The Cauchy–Schwarz inequality states

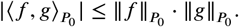

Applying this inequality to Eq. 72 yields the inequality

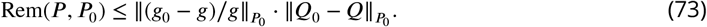

The consistency of TMLE is double-robust in the following sense. Given estimators (*g*_*n*_, *Q*_*n*_) of (*g*_0_, *Q*_0_), *e.g*., constructed using a Super Learner, and assume *g*_*n*_ is a consistent estimator of *g*_0_; the same argument holds if we estimate *Q*_0_ consistently. Consistency implies

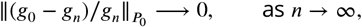

and so 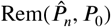 vanishes as *n* → ∞. Then by the definition of Rem(*P*, *P*_0_),

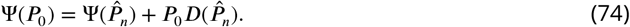

The TMLE step now achieves the following: It updates 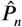 to a final estimate 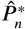, so that

1. 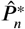 asymptotically has the same *g*-feature and *Q*-feature as *P*_0_, so that it still solves the second-order remainder, 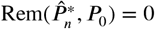, as *n* → ∞; and
2. 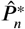 solves the *P*_0_-specific influence curve, *i.e*., 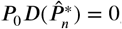, so that 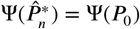.

We include the proof of the proposition for the convenience of the reader.

*Proof*. First, we recall that the influence curve of Ψ_*υ*_(*P*),

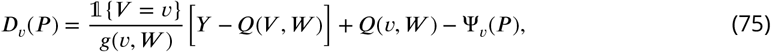

satisfies the property 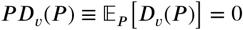, *i.e*., it is mean-centered. Second, using this property, we simplify the second-order remainder Rem(*P*, *P*_0_) as follows:

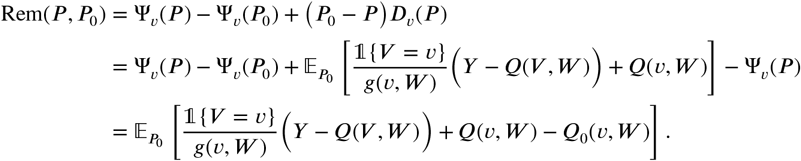

Here, the first equality follows by definition of Rem(*P*, *P*_0_), the second follows by *P D*_*υ*_(*P*) = 0 and 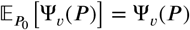, and the third follows by definition of the parameter Ψ_*υ*_(*P*_0_).

Third, we examine the second half of the second-order remainder:

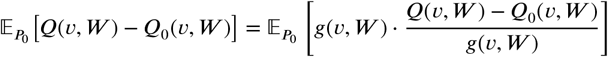

Fourth, we split the remaining term up into two parts,

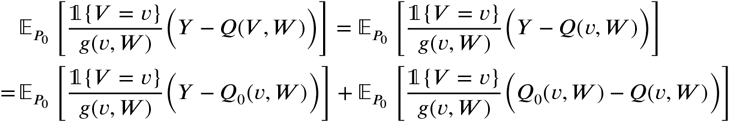

and treat these separately, using the tower rule for both. The first term vanishes:

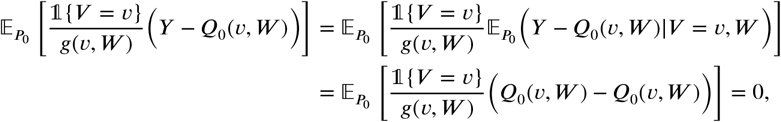

since 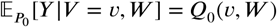. The second term is equal to the following expression:

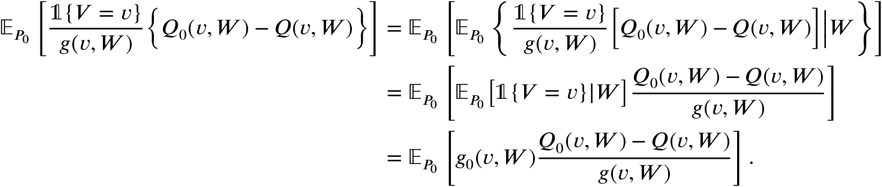

Finally, putting all terms together yields the expression for the second-order remainder:

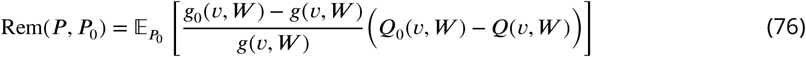

This completes the proof.

## Appendix B Residuals of one fit as dependent variable in another fit

In order to provide a fast method for estimating effect sizes of large numbers of variants on traits, the 2-step residual fitting, originally proposed in GRAMMAR (***Enko et al., 2007***), has been used in the literature to perform GWAS fits. In this procedure, the trait *Y* is first fitted linearly, either using linear regression or an LMM model, as a function of the covariates and random genetic effects. The residual of the fit, *i.e*., *Y* − *Y*_predict_, is then used a dependent variable in a second linear regression model, to estimate the effect size of the variants of interest. However, this procedure results in biased effect estimates and ‘conservative’ tests (***Svishcheva et al., 2012***). Here we demonstrate that the term ‘conservative’ is misleading for two reasons:

1. In the case where a linear model is assumed to be the true model, the estimates of the effect sizes turn out to be less significant than the ground truth value. However, this is only true when *all* relevant covariates have been regressed out in the first step, and *only* the single variant of interest is regressed in the residual fitting step (second step). In contrast, if the residuals from the step 1 are used to obtain the effect size of a variant in the presence of another variable in the fit, the quantities of interest can be under- or over-estimated, and even change sign. This happens, for example, when environmental factors and other variants of interest (*e.g*., for epistasis quantification) are not taken into account in step 1. The extent of under- or -overestimation depends on the degree of dependence (*e.g*., correlation) between the first set and the second set of variables. The issue of 2-step residual fitting has been demonstrated using simulations in other literature, *e.g*., in economics (***Chen et al., 2018***).
2. When the ground truth is non-linear, it is unknown in general what the effect of residual fitting is on the validity of the estimates. These estimates may be under- or over-inflated, or change size.

GRAMMAR-Gamma (***Svishcheva et al., 2012***) introduced a correction factor for the test-statistic and effect size estimates. However, this correction factor only applies when the (unknowable) ground truth is in fact linear and the mistake described in (1) above are not made. If the (unknow-able) ground truth follows a different model, an (unknowable) different analysis is required with (unknowable) new correction factor being applied to fit of the data. For more sophisticated and powerful models, it may be far more complicated to obtain the required correction. Therefore, this is a time-consuming procedure that attempts to treat the symptom instead of the cause of the issue, namely, the 2-step residual fitting procedure. This process should thus be entirely avoided.

**Figure 8.**
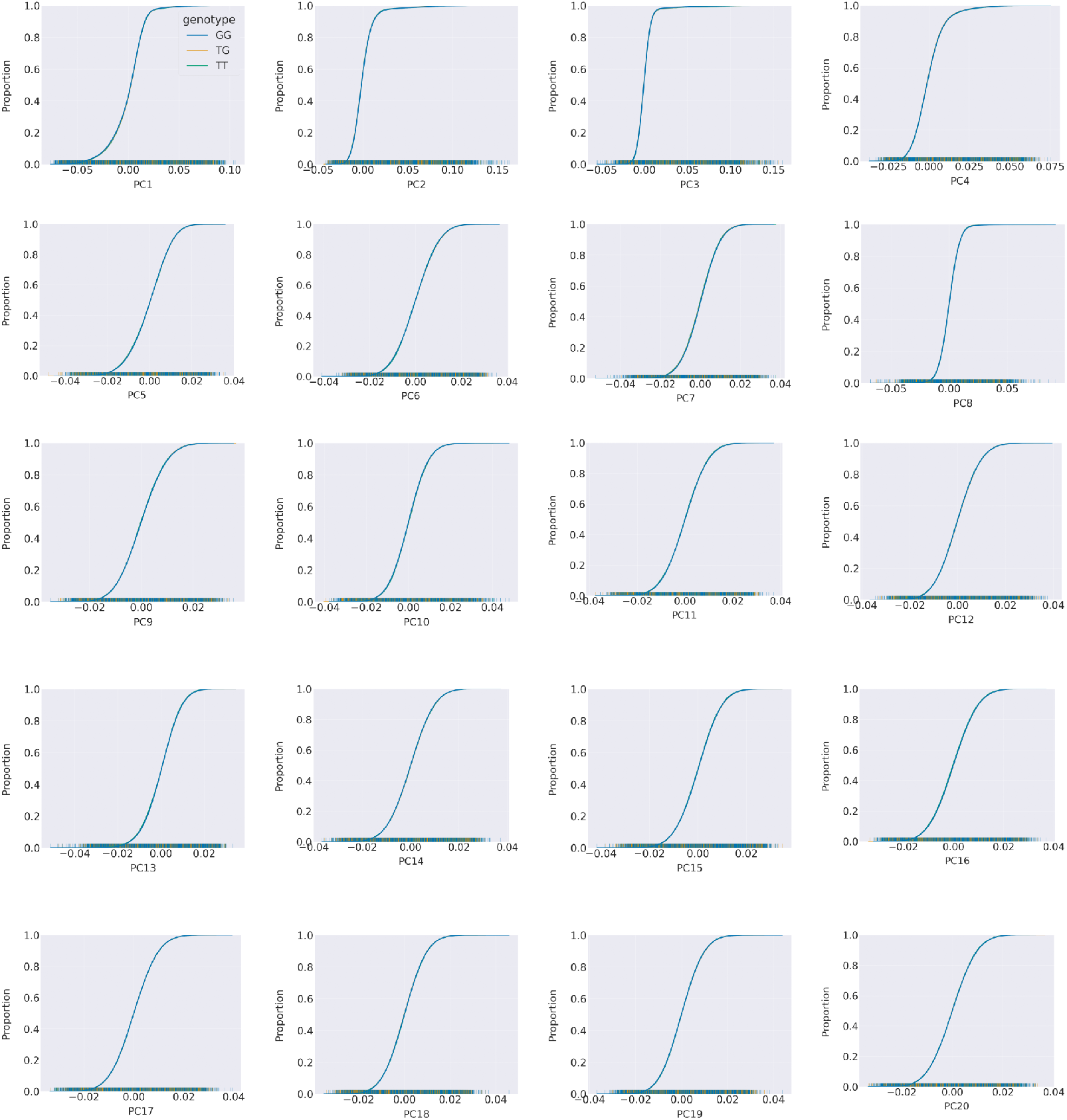
Cumulative frequency diagrams of variant rs1045570 for PCs 1-20.

## Supplementary figures

### Supplement to Figure 5: Population stratification PCA analysis

The figures 8 - 11 are empirical cumulative frequency (ECDF) diagrams.

The aim of these figures is to establish which of the covariates listed by UK Biobank are confounders and which PC number these should take into account. The figures can also be used to establish evidence of population structure based on variants of interest. Large differences between the separate distributions lines indicate a difference between different populations and therefore evidence that population structure is driven by this covariate. The list of variants investigated in this paper is: rs1045570, rs1421085, rs3755967, and rs7971418. Figures 8 - 11 clearly show there is no evidence that population structure is driven by differences in alleles at these variants and therefore, the variants should not have a confounding effect in the analysis. Figure 12 shows the ECDF for self-reported ethnicity. This figure is evidence of population structure in the UK Biobank cohort among participants who self-reported as White ethnicities (including White British, White Irish, White and Any Other White Background). Here we can see that PCs 1-6 have evidence of population structure driven by ethnicity and that this should be included as a confounder in the Super Learner.

**Figure 9.**
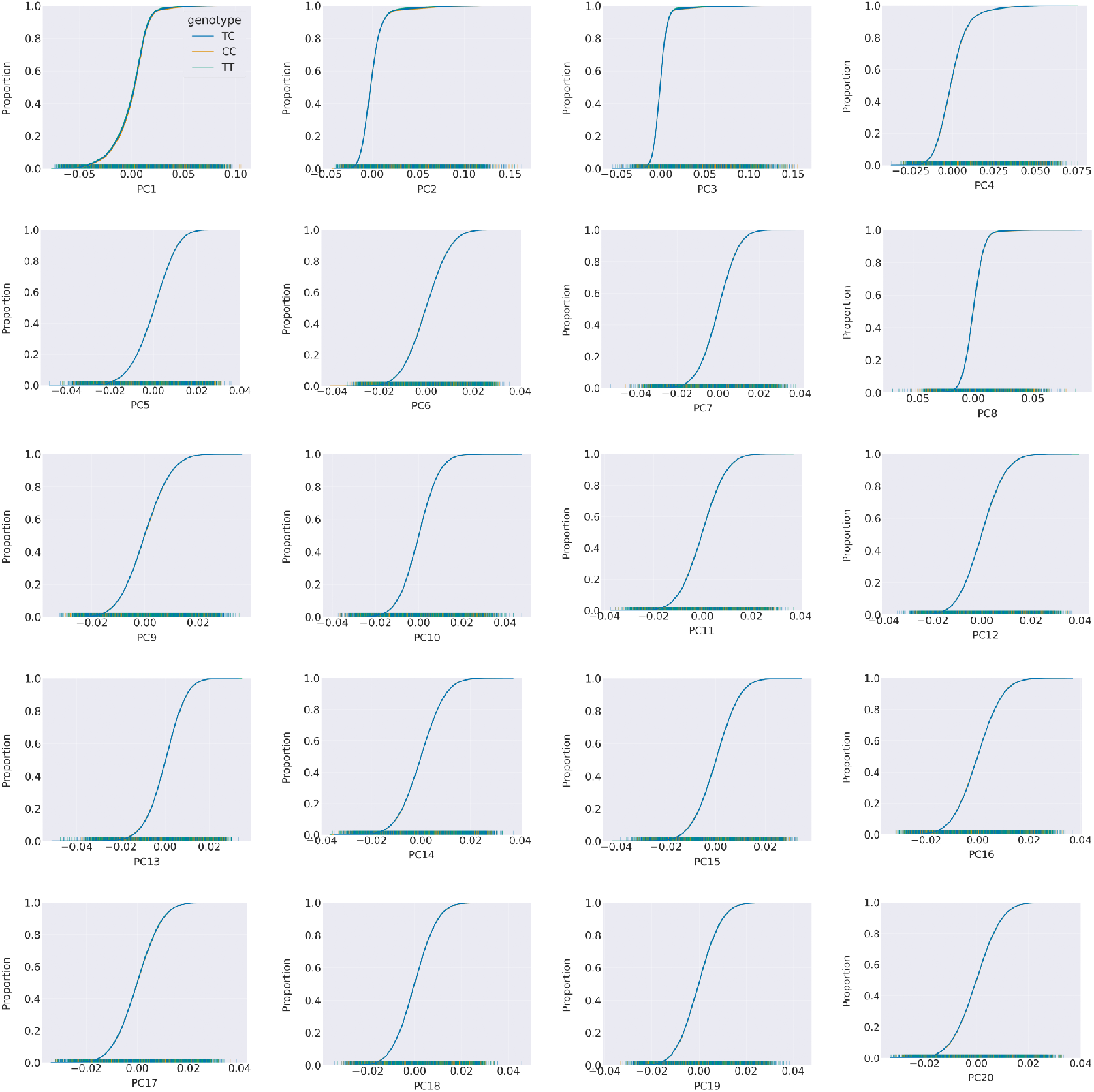
Cumulative frequency diagrams of variant rs1421085for PCs 1-20.

**Figure 10.**
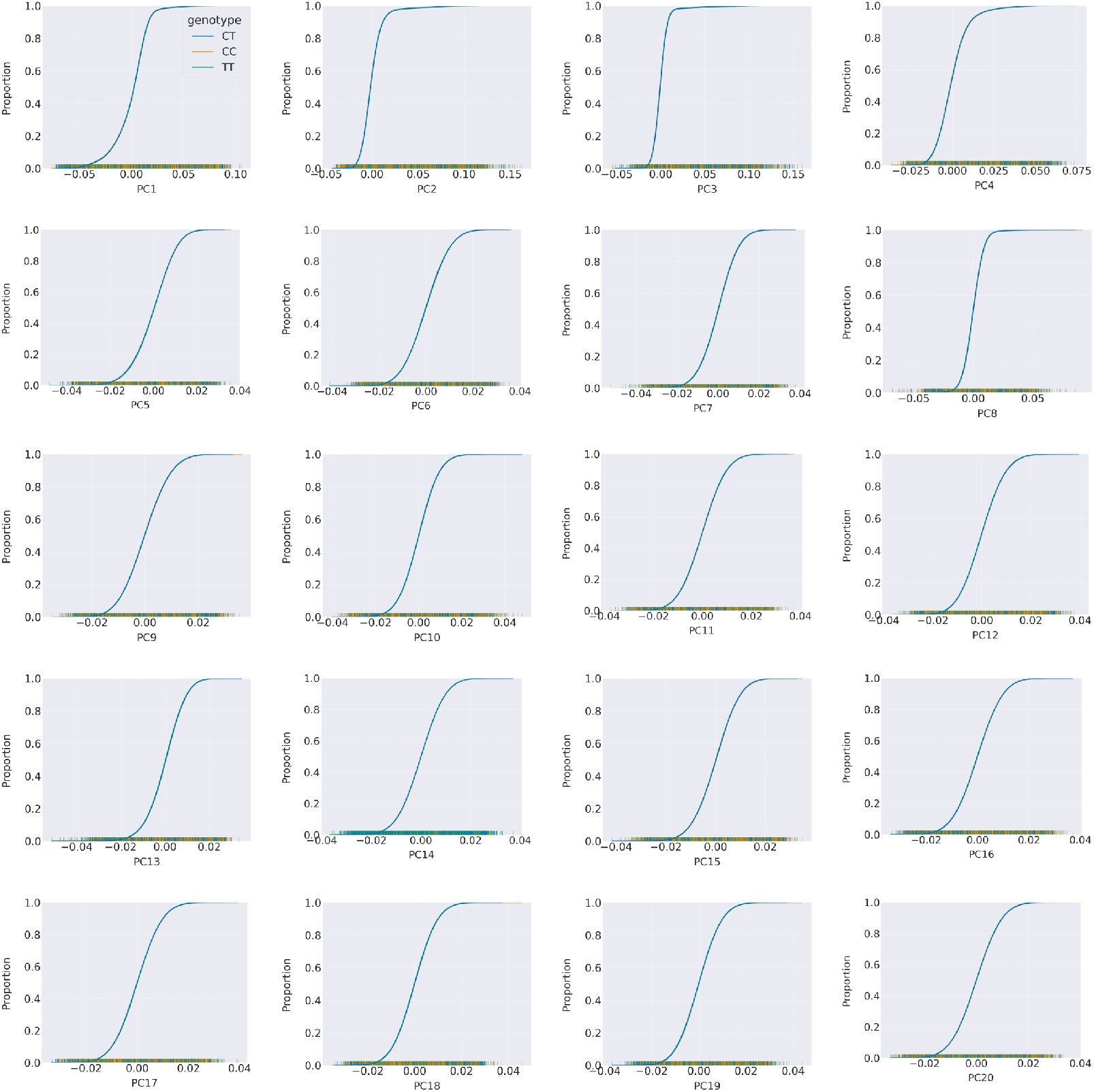
Cumulative frequency diagrams of variant rs3755967 for PCs 1-20

**Figure 11.**
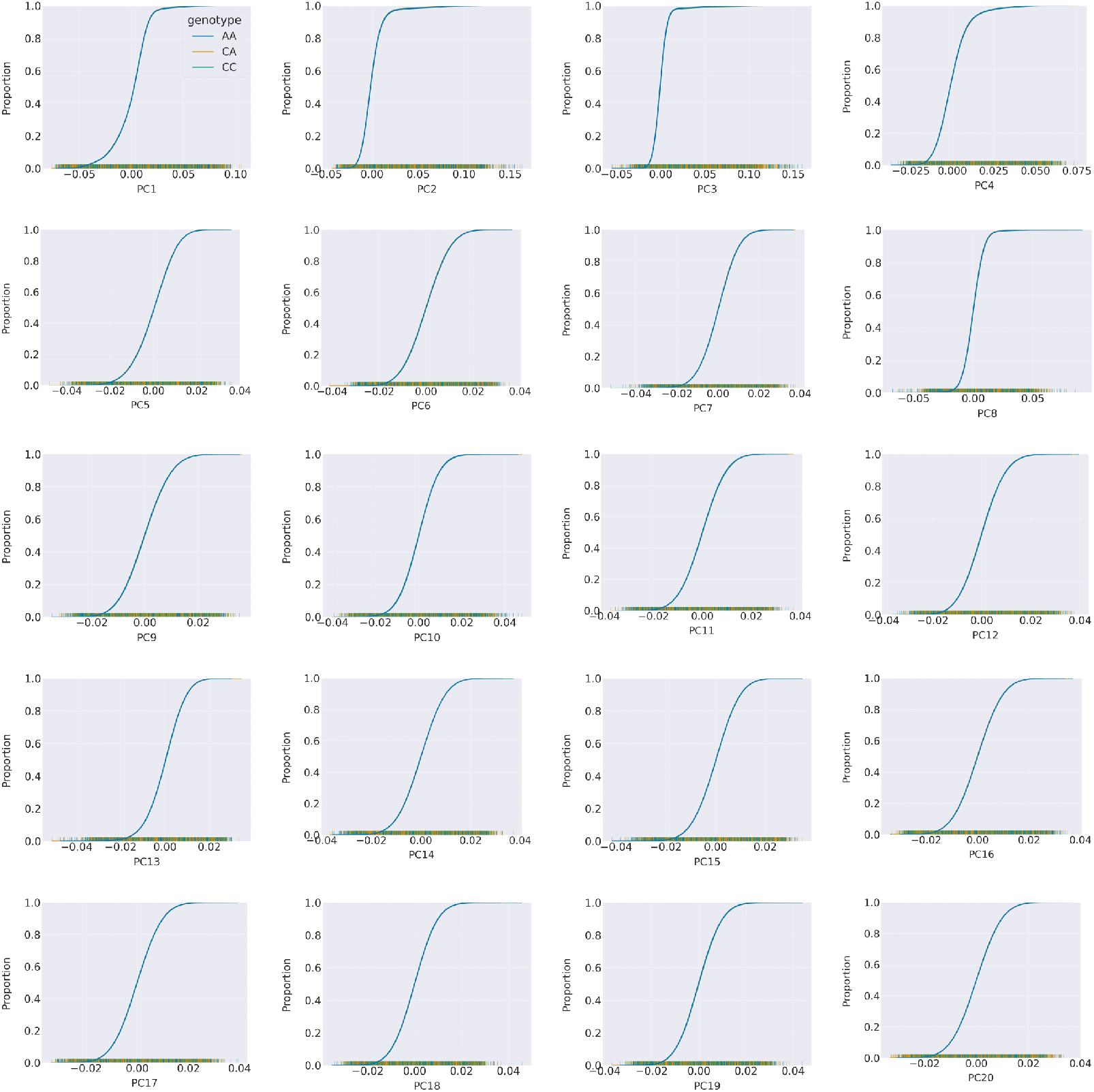
Cumulative frequency diagrams of variant rs7971418 for PCs 1-20

**Figure 12.**
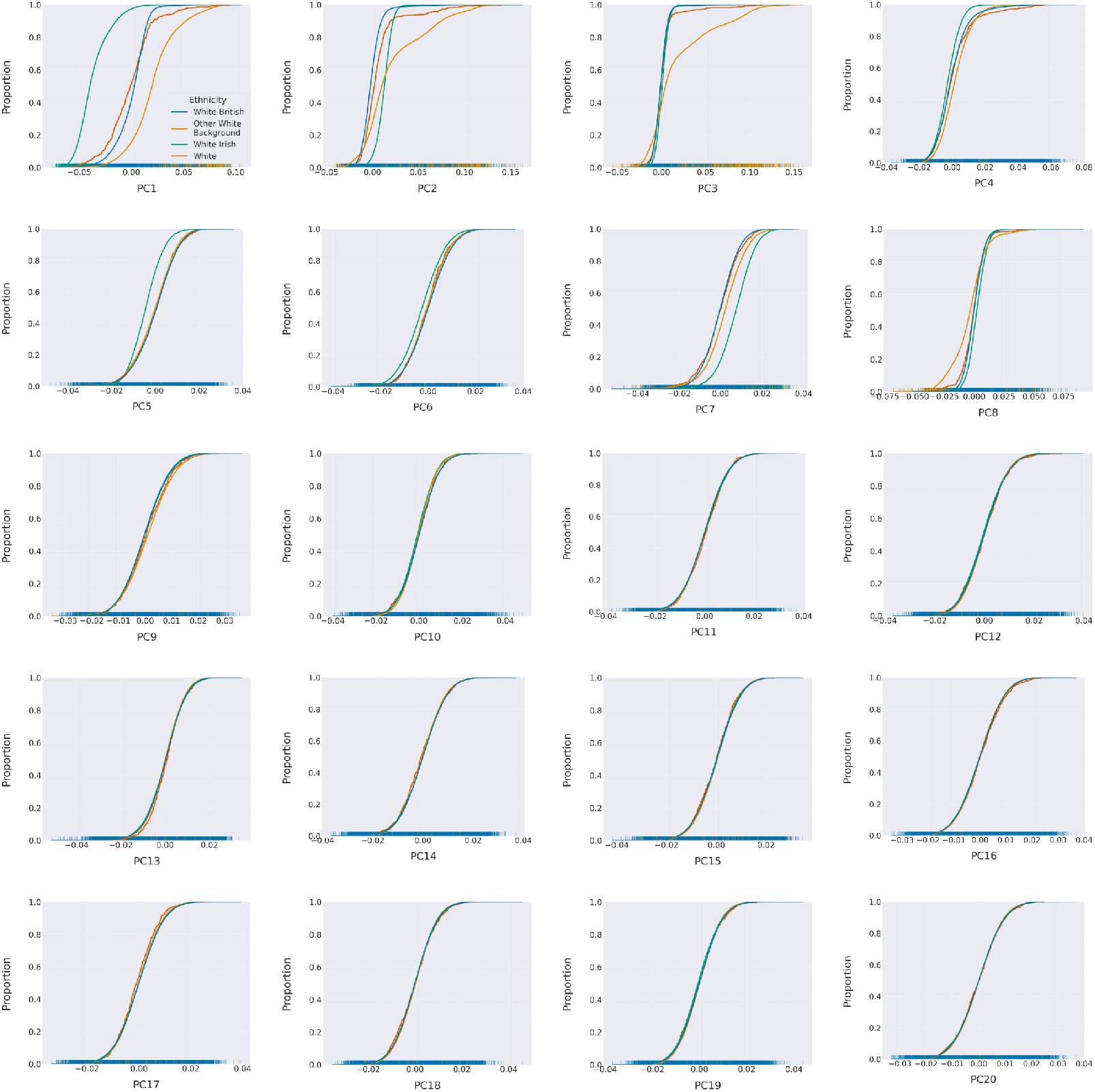
Cumulative frequency diagrams of self-reported ethnicity for PCs 1-20

**Figure 13.**
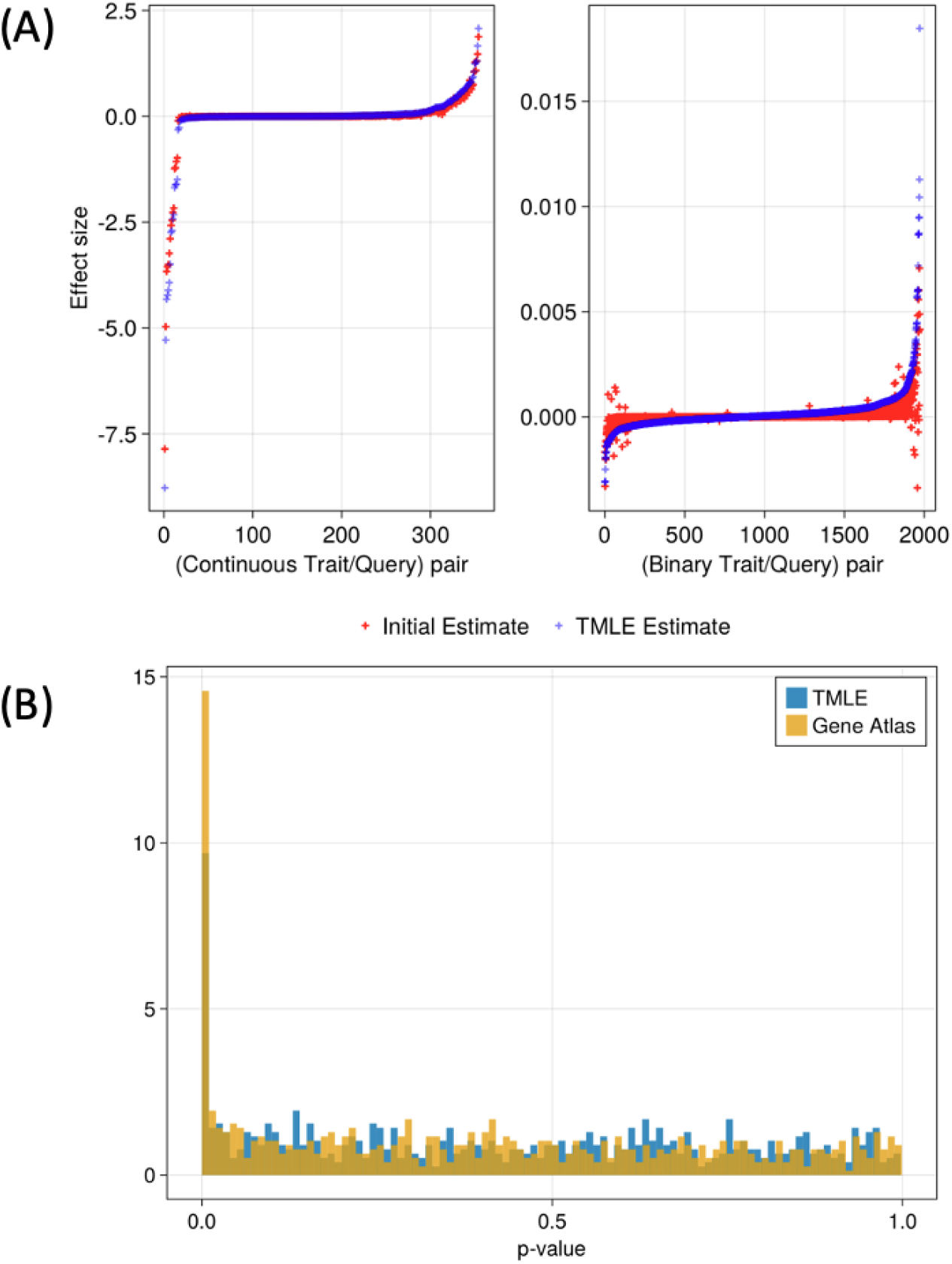
(**A**) Illustration of the difference between the initial estimate, reported by Super Learning, and the TMLE after the targeting step. (**B**) Comparative distribution of the p-values reported by TMLE vs GeneATLAS.

### Supplement to Figure 6: Targeted update of our estimate

We provide additional information regarding the effect of the targeting step on effect sizes and associated p-values. Surprisingly, in most cases, the initial estimate is shifted upward by the TMLE update (see Fig. 13). However, because the variance associated with those estimates is large, they will for the most part not be reported as significant. This behavior is confirmed by the associated p-value comparison, where the distribution of TMLE p-values is slightly shifted towards upper values as compared to GeneATLAS p-values.

In Fig. 14, we present additional information regarding population dependence structure.

**Figure 14.**
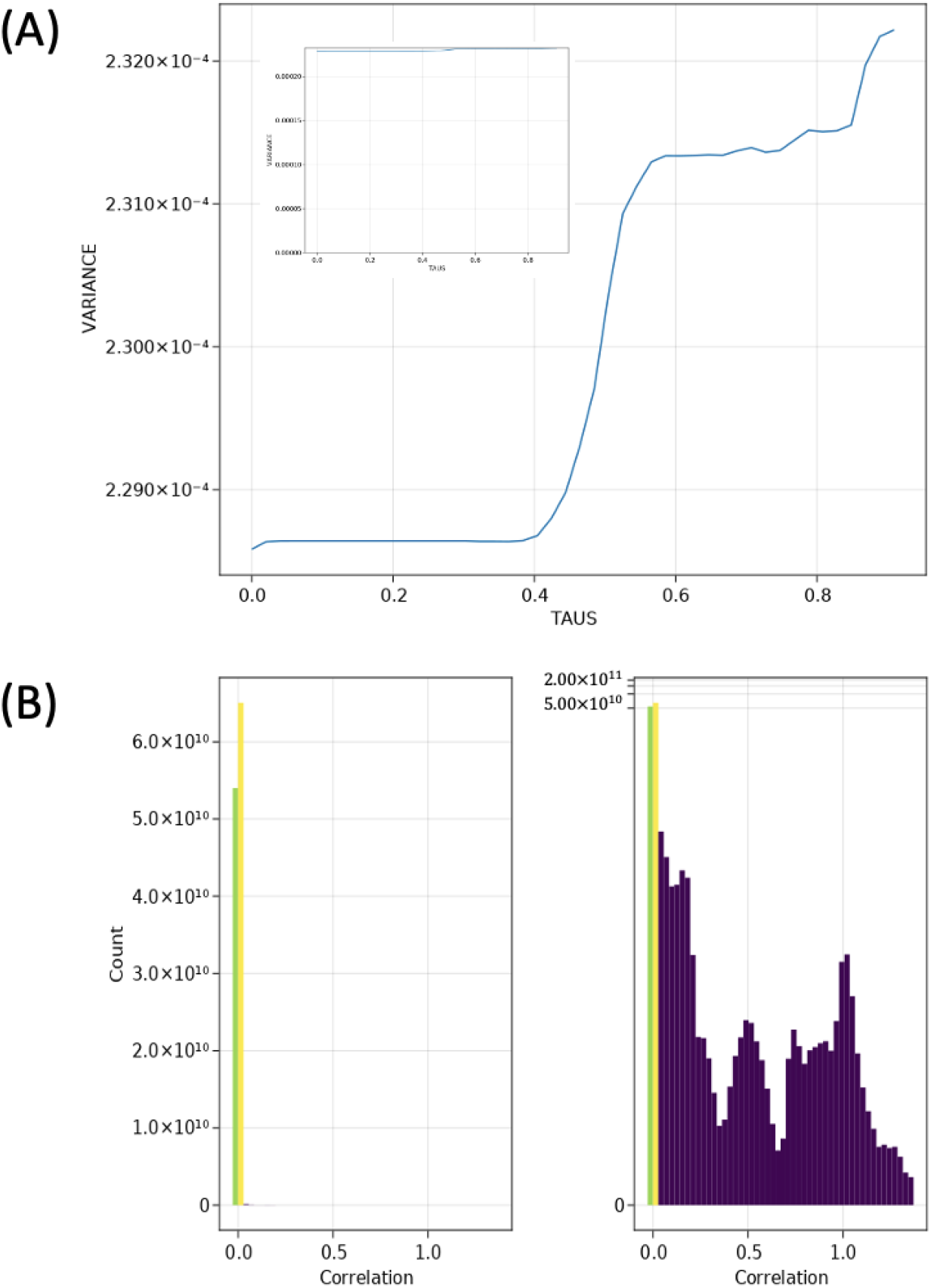
(**A**) A sample curve obtained via the SP estimation method (on two scales). The curve is increasing as we incorporate more dependent individuals in the estimation. (**B**) Histogram of the genetic relationship matrix using both a regular and log10 scale for the full UK Biobank population (488, 376 individuals). As can be seen from the histogram the distribution is highly concentrated around 0. This is indicating that individuals in the UK Biobank are, to a large extent, genetically independent from one another. This may explain why the sieve variance correction has relatively little effect on the variance.

